# Analysis of the high-order conformational changes in glyceraldehyde-3-phosphate dehydrogenase induced by nicotinamide adenine dinucleotide, adenosine triphosphate, and oxidants

**DOI:** 10.1101/2024.11.11.622902

**Authors:** Himari Suzuki, Yuki Nicole Makiyama, Yuta Watanabe, Hideo Akutsu, Michiko Tajiri, Yoko Motoda, Ken-ichi Akagi, Tsuyoshi Konuma, Satoko Akashi, Takahisa Ikegami

**Affiliations:** Graduate School of Medical Life Science, Yokohama City University, 1-7-29 Suehiro, Tsurumi, Yokohama, Kanagawa 230-0045, Japan

**Keywords:** ATP, GAPDH, glyceraldehyde-3-phosphate dehydrogenase, NAD^+^, nicotinamide adenine dinucleotide, nitrosylation, moonlighting protein, oxidation, subunit dissociation

## Abstract

Glyceraldehyde-3-phosphate dehydrogenase (GAPDH) catalyzes the sixth step of glycolysis. As a moonlighting protein, GAPDH interacts with other molecules, seemingly unrelated to glycolysis, to exert additional functions, such as inducing apoptosis. However, specific mechanisms underlying these interactions remain unclear. Therefore, in this study, we aimed to elucidate these mechanisms by analyzing human and porcine GAPDHs based on their three-dimensional (3D) structures using various biophysical techniques, including nuclear magnetic resonance spectroscopy, gel filtration chromatography, and thermal shift assay. Although GAPDH became unstable when nicotinamide adenine dinucleotide (NAD^+^) was removed (*apo* state), the 3D structure of the tetramer was maintained regardless of the temperature. However, in the presence of adenosine triphosphate (ATP), GAPDH split into dimers at low temperatures, exposing the interface between the dimers and increasing the flexibility of the side chains at the site. Moreover, subunit splitting also occurred upon exposure to oxidizing agents, such as hydrogen peroxide. These findings suggest that GAPDH maintains a stable tetramer in the presence of NAD^+^ but becomes unstable and easily oxidized upon NAD^+^ depletion. When multiple residues, including those other than the cysteine residue at the active site, are oxidized by reactive oxygen species or nitric oxide, or when it interacts with ATP, GAPDH splits into dimers. This subunit splitting may trigger interactions with other molecules.

## Introduction

Glyceraldehyde-3-phosphate dehydrogenase (GAPDH) functions in the sixth step of glycolysis, where it reversibly catalyzes the oxidative phosphorylation of glyceraldehyde-3-phosphate to yield 1,3-bisphosphoglycerate while concurrently converting the cofactor, nicotinamide adenine dinucleotide (NAD^+^), to its reduced form, NADH ^1,2^. GAPDH accounts for 5–15% of the soluble proteins in cells ^3^ and is often used as a housekeeping gene in biological experiments. Notably, eukaryotic GAPDHs, under certain circumstances, perform additional functions seemingly unrelated to glycolysis and are known as moonlighting proteins ^3,4^. To perform these additional functions, GAPDHs interact with various other biomolecules, including proteins, nucleic acids ^5^, and heme molecules ^6^. For example, when the active site cysteine residue (Cys152) is nitrosylated (GAPDH-SNO), it interacts with a ubiquitin ligase, Siah1, and the complex subsequently translocates to the nucleus to initiate apoptosis ^7^.

Crystal structures have shown that GAPDHs generally exhibit a homotetramer conformation (4 × 36 kDa), regardless of their origin (Figure 1). However, the conformation of GAPDH when it interacts with other biomolecules to exert its moonlighting functions remains ambiguous owing to the lack of atomic-level structures in such complexes. When interacting with biomolecules, GAPDH may change its quaternary structure to a dimeric or monomeric form, instead of retaining its homotetrameric form. Some reports suggest that the dimeric form of GAPDH interacts with RNA ^8^, tubulin ^9^, and actin ^10^ and that its monomeric form performs uracil-DNA glycosylase functions in the nucleus ^11^.

**Figure 1:**
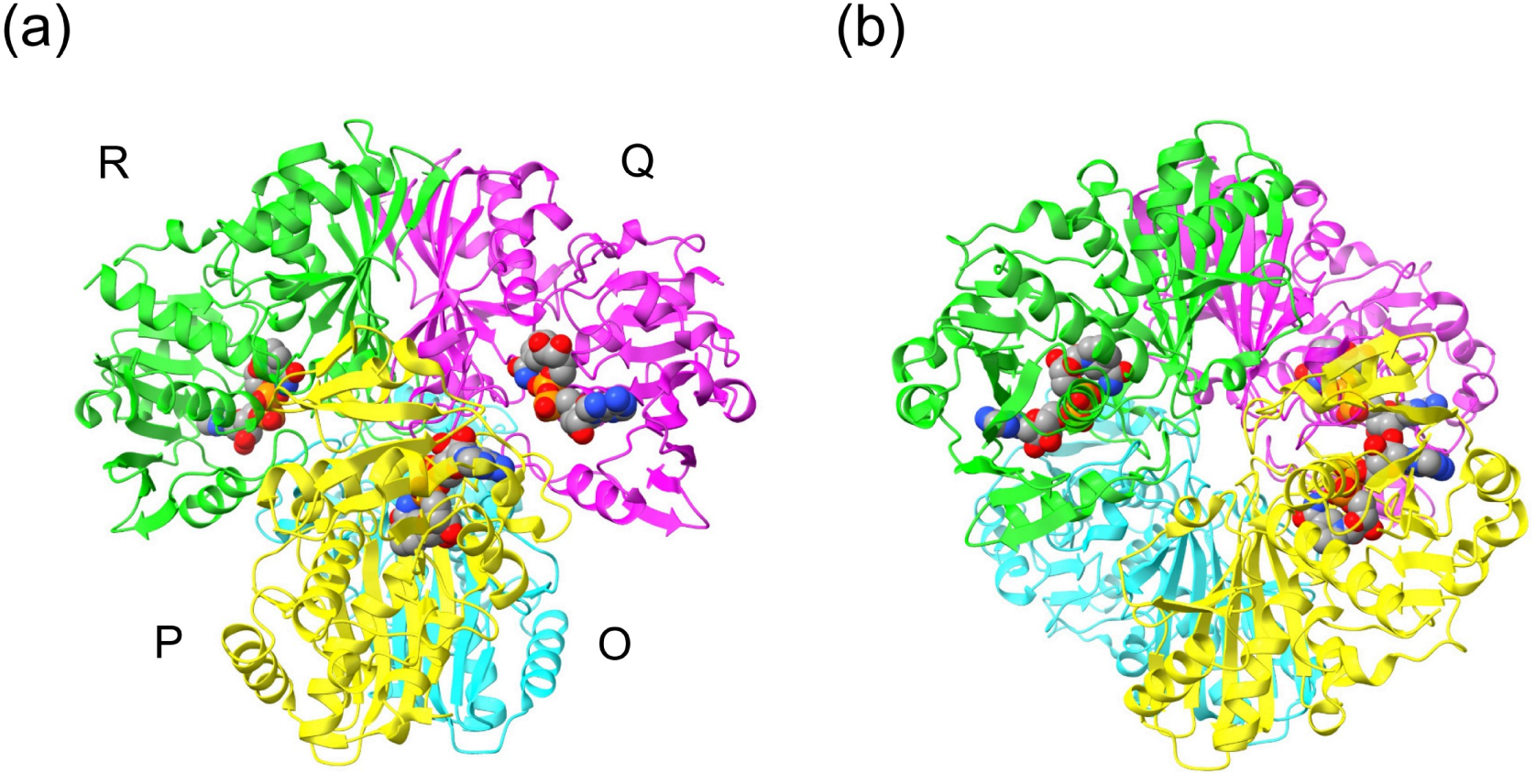
Crystal structure of human GAPDH tetramer. The structure was drawn using ChimeraX ^12^, based on the PDB coordinates from 1u8f ^13^. The subunits are labeled O, P, Q, and R, following the nomenclature in 1u8f. Structure (b) is a 45° rotation of structure (a) around the vertical y-axis. The NAD^+^ molecules attached to the P, Q, and R subunits are represented in a sphere model.

Dissociation of GAPDH subunits and their interactions with other molecules may require chemical modifications, such as nitrosylation and oxidation of cysteine and methionine residues ^14^. However, there are also data showing that GAPDH in the *apo* state, in which no NAD^+^ cofactor is attached, already dissociates into dimers and monomers. Lakatos *et al*. investigated the subunit composition of GAPDH extracted from porcine skeletal muscle at various concentrations via analytical ultracentrifugation ^15^ and found equilibrium states among tetramer, dimer, and monomer *apo* forms ^16^. They also reported that the NAD^+^-saturated *holo* form retained its tetrameric conformation even under conditions where the *apo* form was split (at 5 °C and < 300 µg/mL). A similar phenomenon has been reported for GAPDH extracted from rabbits ^17^. Many variants mimicking chemical modifications have been investigated ^18^. However, it remains unclear whether specific events, such as NAD^+^ dissociation, chemical modification, and induced fit upon binding with other biomolecules, trigger tertiary and quaternary structural changes in mammalian GAPDHs. Additionally, whether such conformational changes lead to functional shifts in GAPDHs is also unclear.

To clarify these points, in this study, we aimed to analyze the conformations of human and porcine GAPDHs (hGAPDH and pGAPDH, respectively) in the *apo* and *holo* states in the presence and absence of additives (adenosine triphosphate [ATP], NAD^+^, and oxidants) using nuclear magnetic resonance (NMR) spectroscopy. NMR spectroscopy provides conformational information at the atomic level for proteins with flexible and transitional structures that are difficult to analyze by X-ray crystallography. We found that most GAPDH molecules in the *apo* state retained a homotetrameric structure similar to that in the *holo* state, even at low temperatures. Consistent with previous reports, the *holo* state was quite stable, whereas the *apo* state aggregated over time. Our findings suggest that transition to the *apo* state via the removal of all NAD^+^ cofactors does not directly cause any significant quaternary structural change but makes GAPDH highly unstable and susceptible to chemical modifications including oxidation, thus promoting subsequent changes in interactions and functions.

## Results

### Almost all GAPDH samples used in the experiments were pretreated with activated charcoal

Unlabeled and isotopically ^15^N/^13^C-labeled hGAPDH (wild-type [wt] and C152S variant hGAPDH) and pGAPDH (wt pGAPDH) were prepared for subsequent analyses. To enhance NMR sensitivity, methyl-selectively labeled GAPDHs were also prepared; only the methyl groups of Met, Ile^δ1^, Leu, and Val residues were labeled with ^1^H/^13^C, whereas the other groups were uniformly labeled with ^2^H, ^15^N, and ^12^C ^19^. From genetically recombinant *Escherichia coli* (*E. coli*) cells cultured in 100 mL of D_2_O-based M9 minimal medium, purification of methyl ^1^H/^13^C labeled GAPDHs, otherwise deuterated, yielded approximately 40−200 µg, corresponding to a subunit concentration of 4−20 µM for NMR analyses. Despite induction at 15 °C, approximately 70% of the expressed proteins formed inclusion bodies in *E. coli* cells. When a plasmid construct containing the *hgapdh* gene lacking an N-terminal (His)_6_-tag was used, approximately half of the expressed proteins dissolved; however, insufficient purification often resulted in non-specific cleavage by contaminating proteases. Unless otherwise specified, all hGAPDHs and pGAPDHs were treated with activated charcoal before analysis, resulting in approximately half of the purified GAPDHs being lost owing to adsorption onto activated charcoal. Then, specific activity of pGAPDH treated with activated charcoal was found to be 102.0 ± 3.9 µmol/min/mg, similar to the reported value (100 µmol/min/mg) for GAPDH extracted from rabbit muscles ^20^. Furthermore, freezing at –80 °C for more than six months did not diminish the enzyme activity, as thawed hGAPDH and pGAPDH exhibited activities of 108.3 and 127.7 µmol/min/mg, respectively. Notably, no significant differences were observed in the activities of GAPDHs with and without the (His)_6_-tag.

### The *apo* form of hGAPDH, detected by electrospray ionization mass spectrometry (ESI-MS), maintained its tetrameric conformation

Using ESI-MS under native-like conditions, we investigated the mechanism by which treatment with activated charcoal altered the number of NAD^+^ cofactors bound to homotetrameric hGAPDH. Without prior treatment with activated charcoal, hGAPDH in 10 mM ammonium acetate exhibited attachment of two, three, and four NAD^+^ cofactors to the GAPDH tetramer (Figure 2a). In contrast, the activated charcoal-treated sample showed unsplit peaks, indicating the presence of the *apo* form with no NAD^+^ bound (Figure 2b). In both cases, no peaks corresponding to species other than homotetramers, such as dimers or monomers, were detected. The tetramer peaks observed in the mass spectra were somewhat broadened, despite examining various measurement conditions. This suggests that some solvent and/or salt may still have been retained at the subunit interface. These data indicate that hGAPDH, at least under the MS measurement conditions, maintains a homotetrameric conformation, even in the *apo* state.

**Figure 2:**
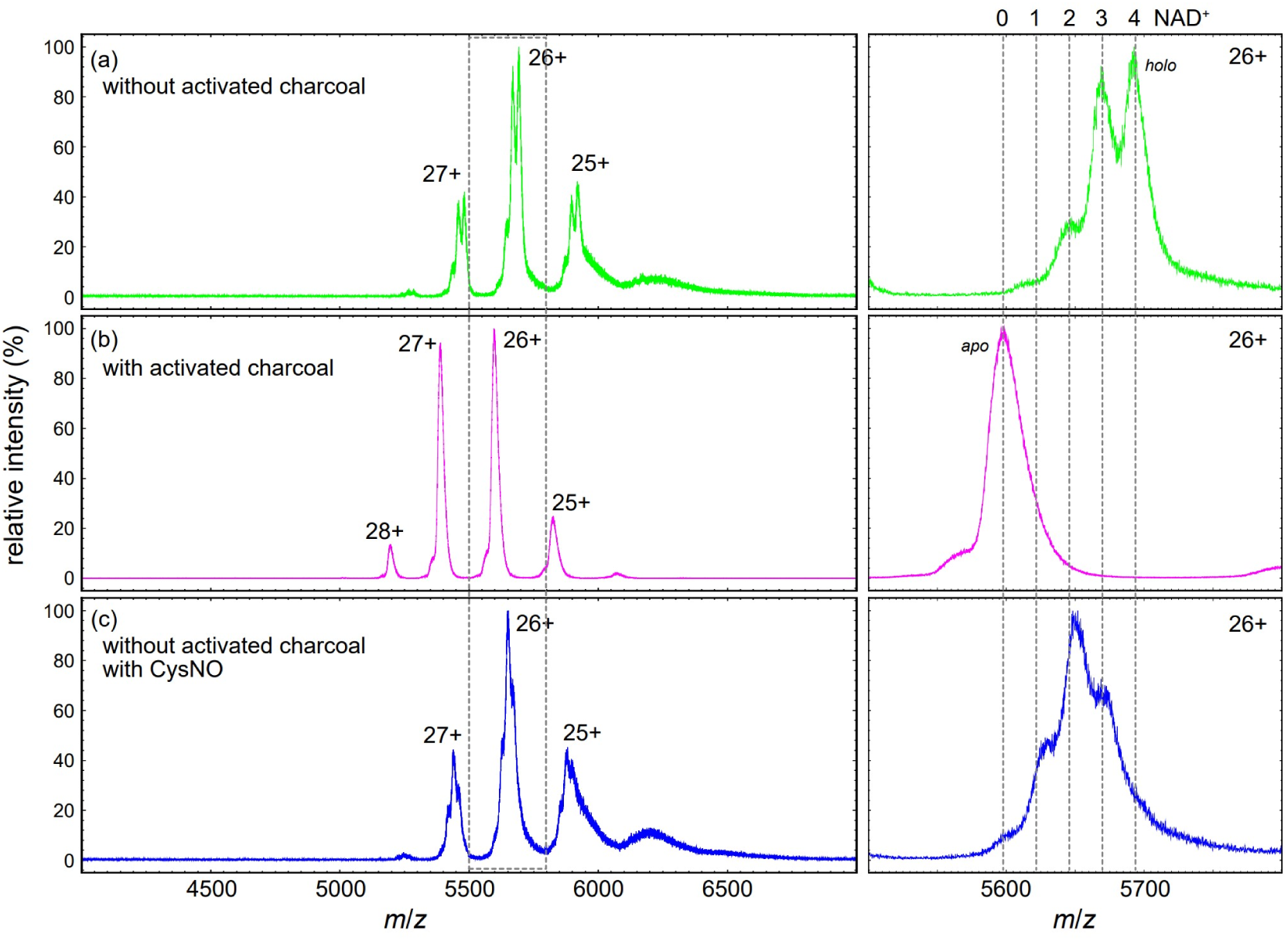
Electrospray ionization-mass spectrometry (ESI-MS) spectra of human glyceraldehyde-3-phosphate dehydrogenase (hGAPDH), obtained under native-like conditions. Spectra were recorded before (a) and after (b) treatment with activated charcoal. (c) Spectrum was acquired after reaction with S-nitrosocysteine (CysNO), without activated charcoal treatment. In the absence of activated charcoal and CysNO treatment (a), the hGAPDH tetramer showed attachment of 1–4 NAD^+^ molecules, whereas after CysNO treatment (c), the tetramer was associated with 0–3 NAD^+^ molecules. Enlarged views of the peak regions corresponding to the 26+ charge state, outlined by dotted lines, are shown to the right of each panel. The numbers of NAD^+^ molecules bound to the homotetrameric form of hGAPDH are also indicated at the top.

### Similar NMR spectra of the *apo* and *holo* forms of hGAPDH indicated that NAD^+^ removal caused no major structural changes

NMR peaks were clearly observed in the two-dimensional (2D) ^1^H-^13^C methyl-transverse relaxation-optimized spectroscopy (TROSY) heteronuclear multiple quantum correlation (HMQC) spectrum ^21^ measured at 40 °C for hGAPDH, even in the *apo* state, treated with activated charcoal immediately before measurement. The peak positions representing the chemical shifts resembled those observed in the spectrum of the *holo* form (Figure 3). Although some peaks exhibited significant differences, these variations likely stemmed from the nuclei situated near the NAD^+^-binding site. The chemical shift changes were thus probably due to localized and minor conformational alterations around the NAD^+^-binding site or the direct influence of NAD^+^. Therefore, the *apo* form did not undergo substantial structural changes, such as subunit splitting, under the tested conditions.

**Figure 3:**
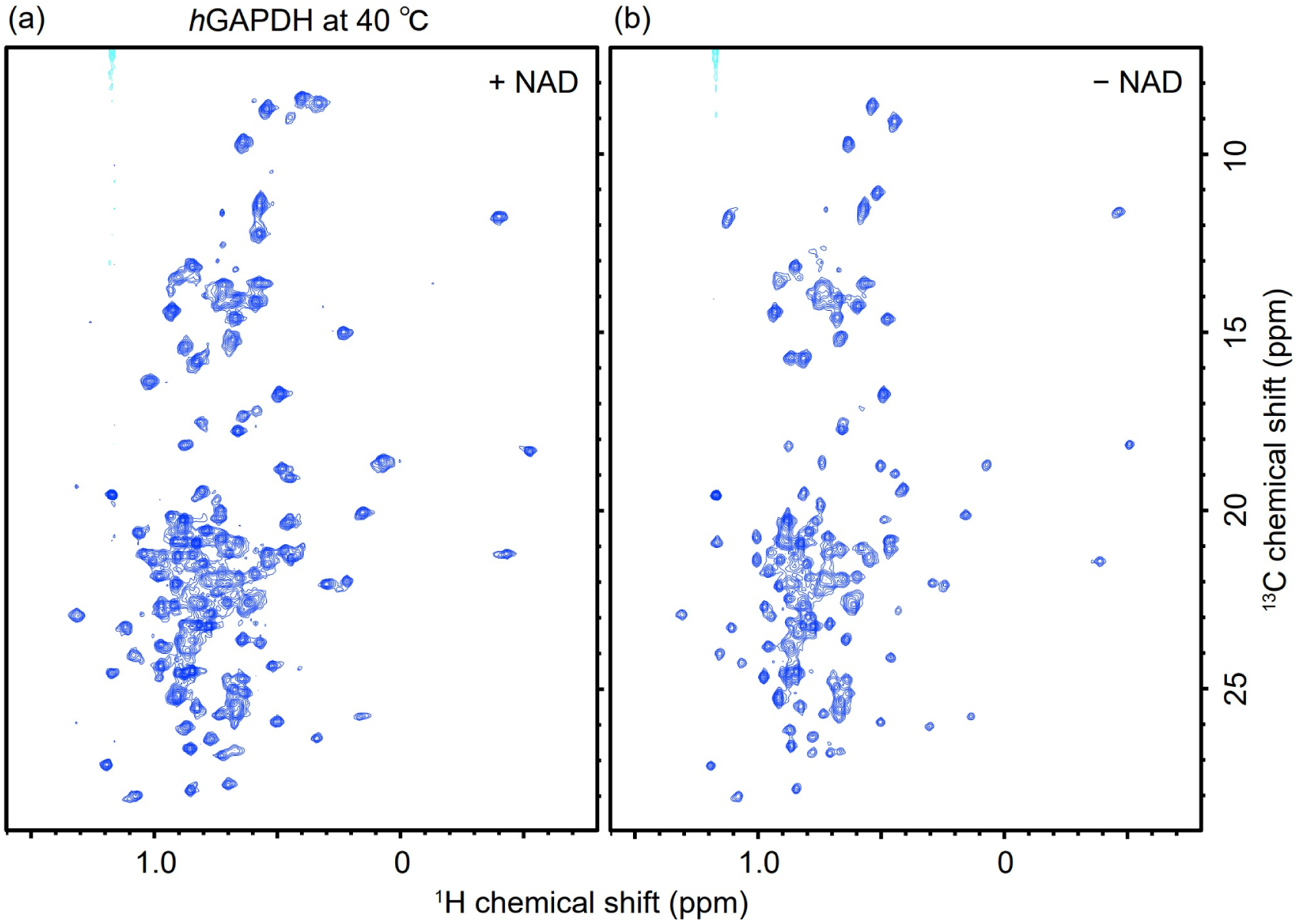
Two-dimensional (2D) ^1^H-^13^C methyl-transverse relaxation-optimized spectroscopy (TROSY) heteronuclear multiple quantum correlation (HMQC) spectra of 4.0 µM {*u*-[^2^H, ^15^N]; Ile^δ^^1^-[^13^CH_3_]; Met^ε^-[^13^CH_3_]; Leu^δ^, Val^γ^-[^13^CH_3_, ^12^CD_3_]}-hGAPDH dissolved in 20 mM sodium phosphate buffer (pH 7.4) containing 10 mM NaCl, 1 mM EDTA, 1 mM dithiothreitol (DTT), and 10% (*v*/*v*) D_2_O in the presence (a) and absence (b) of 2 mM NAD^+^. These spectra were obtained at ^1^H resonance frequency of 950 MHz at 40 °C.

### The *apo* and *holo* forms of pGAPDH also exhibited similar structures

As described above, removal of NAD^+^ caused no significant conformational changes in hGAPDH, at least at room temperature (15–40 °C). In contrast, many studies have focused on the subunit splitting of *apo* GAPDHs, specifically those from yeasts ^22^, rabbits ^17,23^, and pigs ^15^ rather than those from humans. The sequences of rabbit GAPDH and pGAPDH showed similarities, with a multiple sequence alignment (CLUSTAL 2.1) score of 98.8, whereas both sequences differed slightly from that of hGAPDH, with a score of 95.5 (Figure 4). Therefore, we expressed pGAPDH in *E. coli* to examine the possible effects of NAD^+^ on the structure of pGAPDH using NMR. As shown in Figure 5, 2D ^1^H-^13^C heteronuclear single quantum coherence (HSQC) spectra of the *apo* and *holo* forms showed no significant differences in chemical shifts or linewidths. Similar to that observed for hGAPDH, no major conformational changes, such as subunit dissociation, were observed in pGAPDH after removal of NAD^+^, at least at 25 °C.

**Figure 4:**
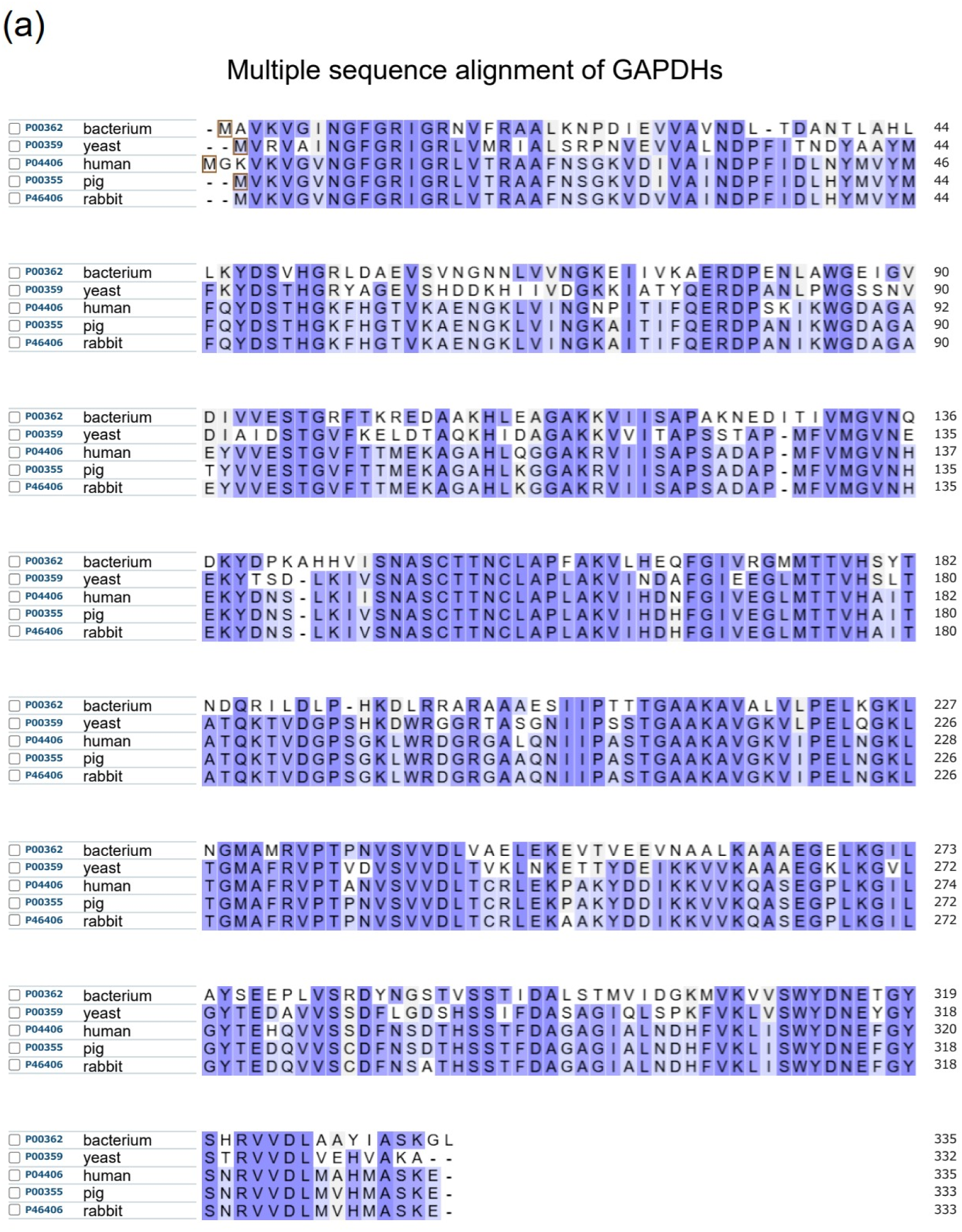
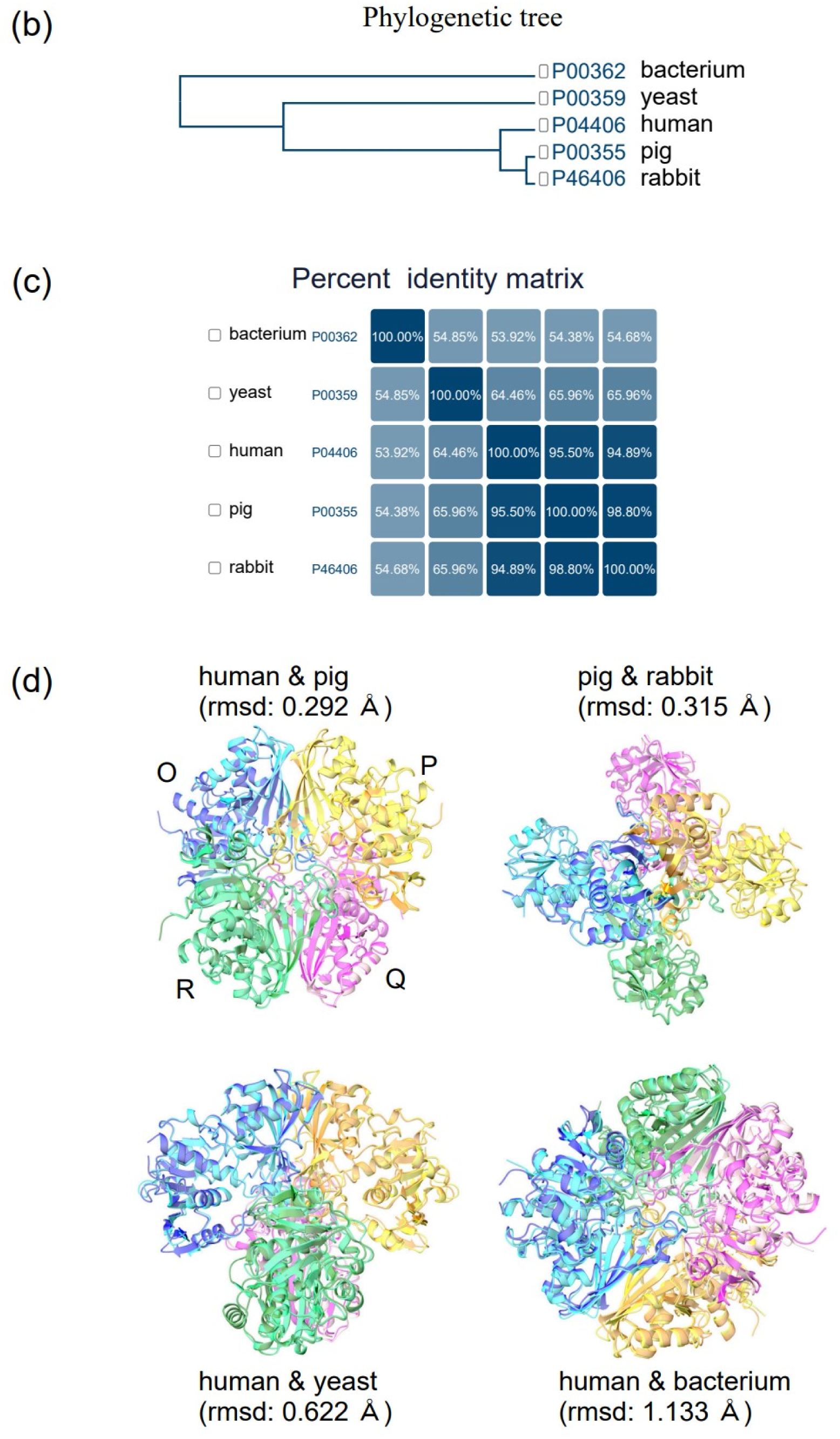
Comparison of GAPDHs derived from different organisms. (a) The amino acid sequences of GAPDHs from human *Homo sapiens* (P04406, G3P_HUMAN), pig *Sus scrofa* (P00355, G3P_PIG), rabbit *Oryctolagus cuniculus* (P46406, G3P_RABIT), yeast *Saccharomyces cerevisiae* (P00359, G3P3_YEAST), and bacterium *Bacillus stearothermophilus* (P00362, G3P_GEOSE) were obtained from the UniProt database (accession numbers shown in parentheses) and aligned using the Align tool in UniProt (Clustal Omega 1.2.4). The degree of amino acid similarity is indicated by the intensity in blue. The accompanying outputs, including (b) a phylogenetic tree and (c) a percentage identity matrix, are also shown. (d) Based on the amino acid sequences, the respective homotetrameric structures were predicted using AlphaFold3 ^24^ and pairwise structural alignments were calculated using ChimeraX ^12^. The root-mean-square deviations (RMSDs) between the C^α^ atoms are also shown. Subunit labels—O (blue and cyan), P (orange and yellow), Q (magenta and pink), and R (green and light green)—are indicated. In the notation “A & B”, “A” corresponds to the structure in dark colors (blue, orange, magenta, and green), and “B” corresponds to the structure in light colors (cyan, yellow, pink, and light green).

**Figure 5:**
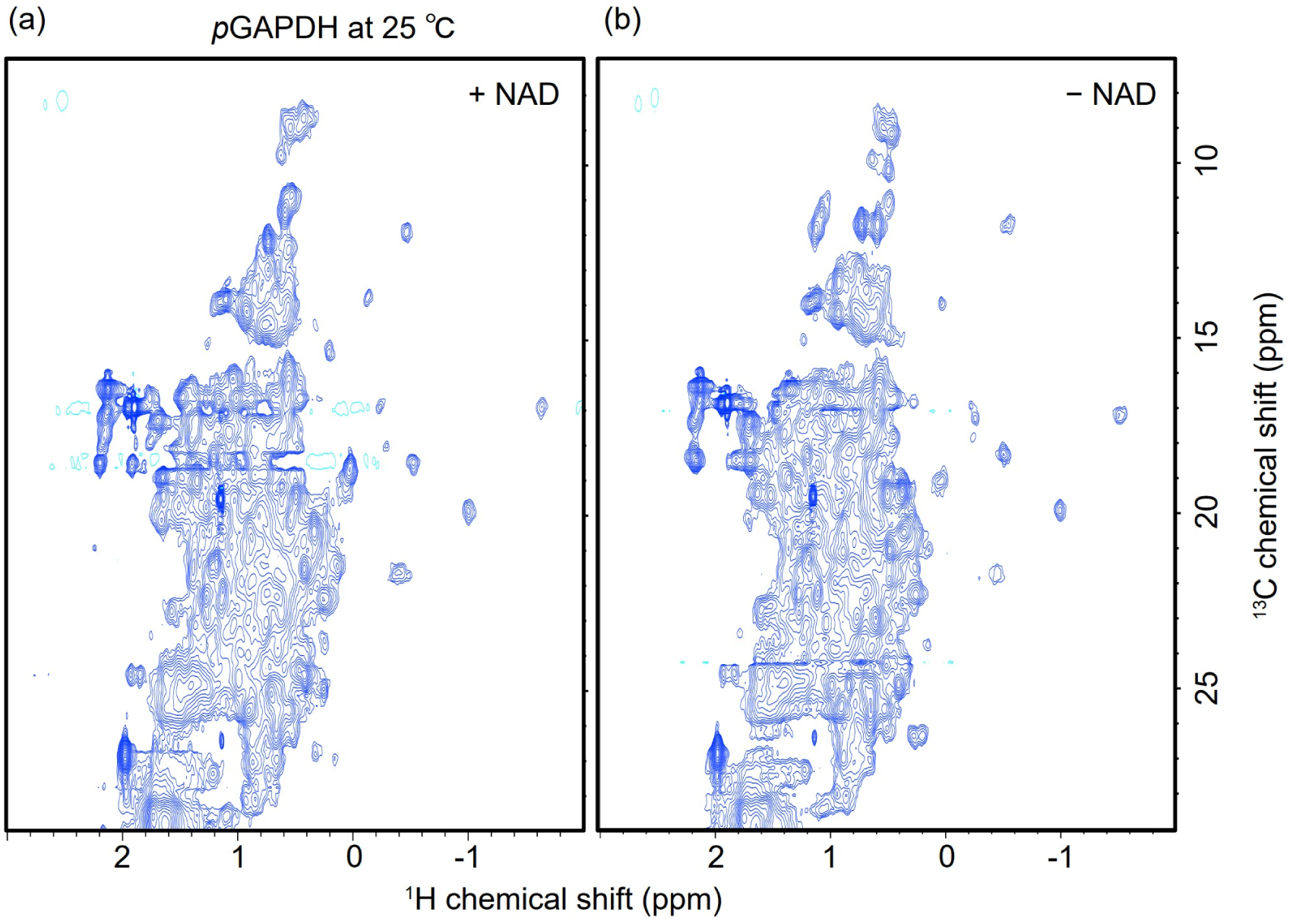
2D ^1^H-^13^C heteronuclear single quantum coherence (HSQC) spectra of *holo*-(a) and *apo*-(b) forms of [^15^N, ^13^C]-porcine GAPDH (pGAPDH). The *holo*-form sample contained 10 mM of NAD^+^. For each sample, the protein was dissolved at a subunit concentration of 100 µM in 20 mM sodium phosphate buffer (pH 7.4) containing 10 mM NaCl, 1 mM DTT, 1 mM EDTA, and 10% D_2_O. The spectra were measured at 25 °C using an 800 MHz NMR spectrometer. Its peaks were broader than those of hGAPDH, as pGAPDH was not deuterated.

### Most *apo* pGAPDH molecules maintained a tetrameric conformation regardless of temperature and concentration

pGAPDHs in the *holo* and *apo* states were subjected to analytical gel filtration chromatography at 4 °C and 28 °C. Both pGAPDHs were eluted at positions estimated to have a molecular weight of 120 kDa, similar to that of the tetramer (144 kDa), regardless of temperature (Figure 6). Even when the concentration of *apo* pGAPDH was reduced from 25 µM to 1 µM, the elution position remained unchanged even at 4 °C. The actual concentration of pGAPDH during column traversal was likely to be below 1 µM. The slightly delayed elution compared with the position estimated from the ideal molecular weight was probably due to a subtle interaction between pGAPDH and the resin. These results indicate that *apo* pGAPDH maintains its homotetrameric conformation. However, in the analysis of the *apo* form at 4 °C, the elution peak of the tetramer was often followed by a smaller peak (marked with an asterisk in Figure 6) at 60 kDa, which was close to the molecular weight of the dimer (72 kDa). In some cases, the two peaks fused, forming a small shoulder adjacent to the tetramer peak. This outcome suggests that, while most *apo* pGAPDH molecules maintain their tetrameric structure at 4 °C, a few populations—too small to be observed by NMR—split into dimers. We performed a similar analysis to determine the molecular weight of hGAPDH by analytical gel filtration. Similar to pGAPDH, hGAPDH eluted at the tetramer position regardless of the presence or absence of NAD^+^ at different concentrations (Figure 7). However, unlike pGAPDH, no small peaks were observed, indicating the absence of dimers. This discrepancy may be attributed to the greater instability of pGAPDH compared to that of hGAPDH, as described below.

**Figure 6:**
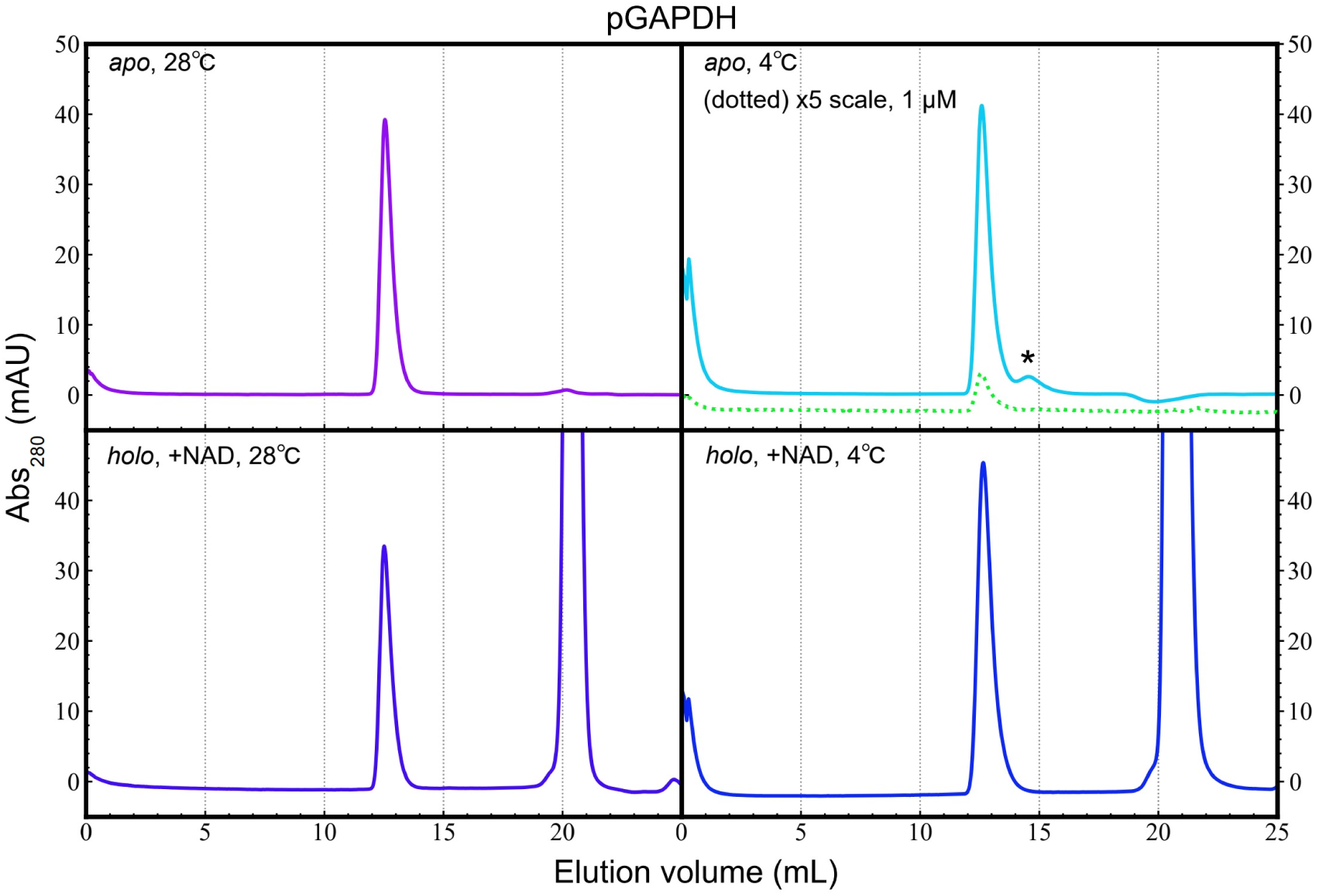
Analytical gel filtration of pGAPDH. Samples (25 µM; 300 µL) with (*holo*) and without (*apo*) NAD^+^ were separately applied to an analytical gel filtration column (Superdex 200 increase 10/300 GL) at 4 and 28 °C, as indicated in each chromatograph, at a flow rate of 0.38 mL/min. The running buffer comprised 20 mM Tris-HCl (pH 8.0) and 200 mM NaCl. For *holo* sample analysis, the injected sample and running buffer contained 2 and 1 mM NAD^+^, respectively. NAD^+^ was eluted at approximately 21 mL volume. The peak indicated by an asterisk possibly originated from a homodimeric component present at a low molar ratio. Additionally, the dotted line illustrates the 5-fold magnified elution curve observed with 1 µM *apo* pGAPDH.

**Figure 7:**
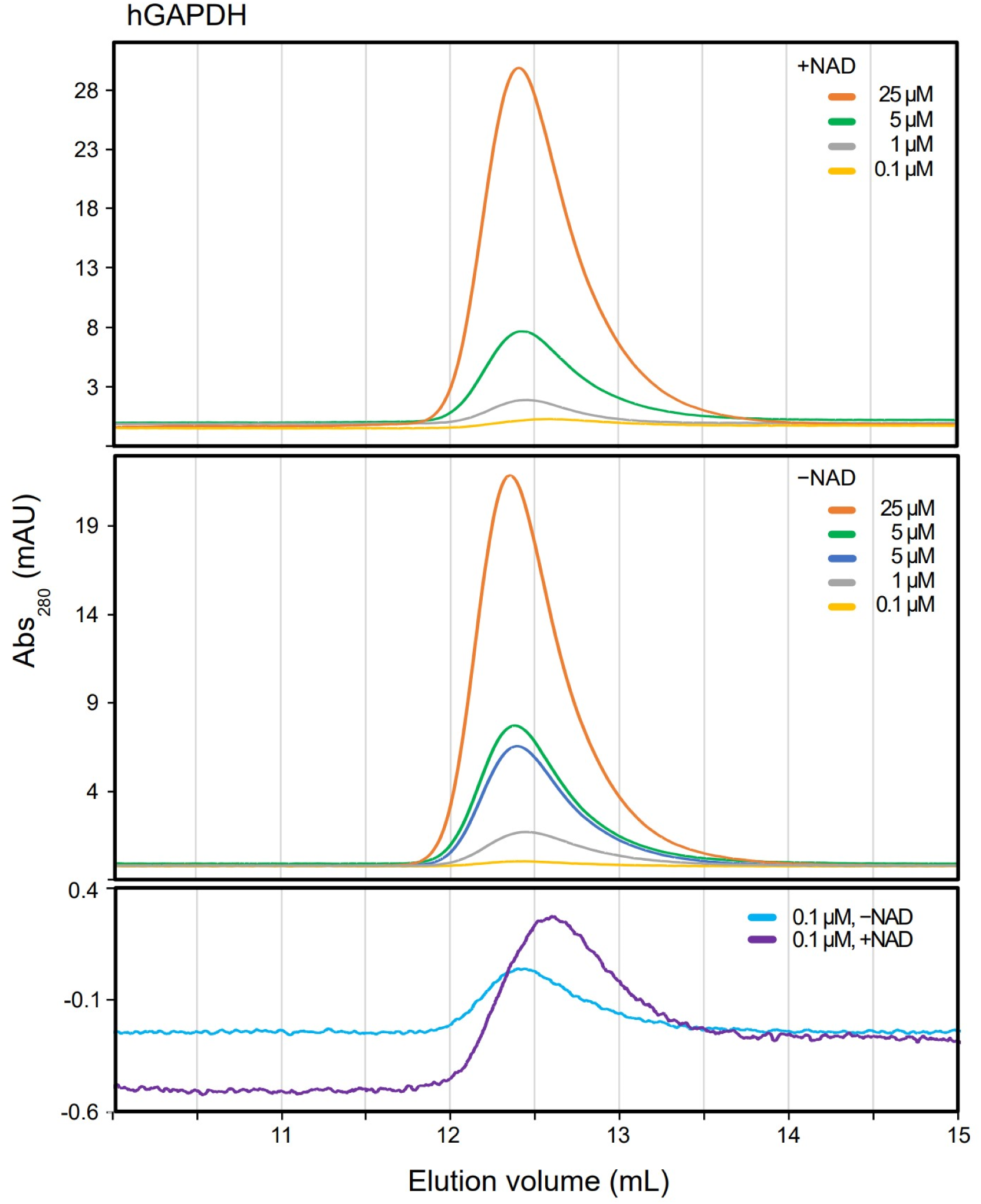
Analytical gel filtration of hGAPDH. (a) *Holo* hGAPDH samples containing 2 mM NAD^+^ were individually loaded onto an analytical gel filtration column (Superdex 200 increase 10/300 GL) at concentrations of 25.0, 5.0, 1.0, and 0.1 µM (each 300 µL) at 5 °C and flow rate of 0.38 mL/min. The running buffer consisted of 20 mM Tris-HCl (pH 8.0), 200 mM NaCl, and 1 mM NAD^+^. Elution volumes at the peak positions (corresponding molecular weights estimated from the calibration curve) were 12.41 mL (130 kDa), 12.43 mL (129 kDa), 12.46 mL (128 kDa), and 12.59 mL (120 kDa), respectively. (b) After removing NAD^+^, *apo* hGAPDH samples were subjected to the same procedure as described in (a), but the solvent or running buffer did not contain NAD^+^. The elution volumes (corresponding molecular weights) were 12.35 mL (134 kDa), 12.37 mL (133 kDa), 12.39 mL (132 kDa), 12.44 mL (129 kDa), and 12.39 mL (132 kDa), respectively. Panel (c) shows an overlay of the expanded elution profiles obtained from (a) and (b), when *apo* and *holo* hGAPDHs (each 0.1 µM) were applied.

### The *holo* form remained stable even at high temperatures, whereas the *apo* form was unstable

Following a 23.5-h NMR measurement at 40 °C, we observed severe precipitation of *apo* hGAPDH, with the peak intensity dropping on average to 40% of that of *holo* hGAPDH. As shown in Figure 3b, precipitation and aggregation notably reduced the peak intensity of the *apo* form. Conversely, *holo* hGAPDH exhibited exceptional stability, maintaining an almost identical 2D spectrum even after storage at 4 °C for 10 months (Figure 8).

**Figure 8:**
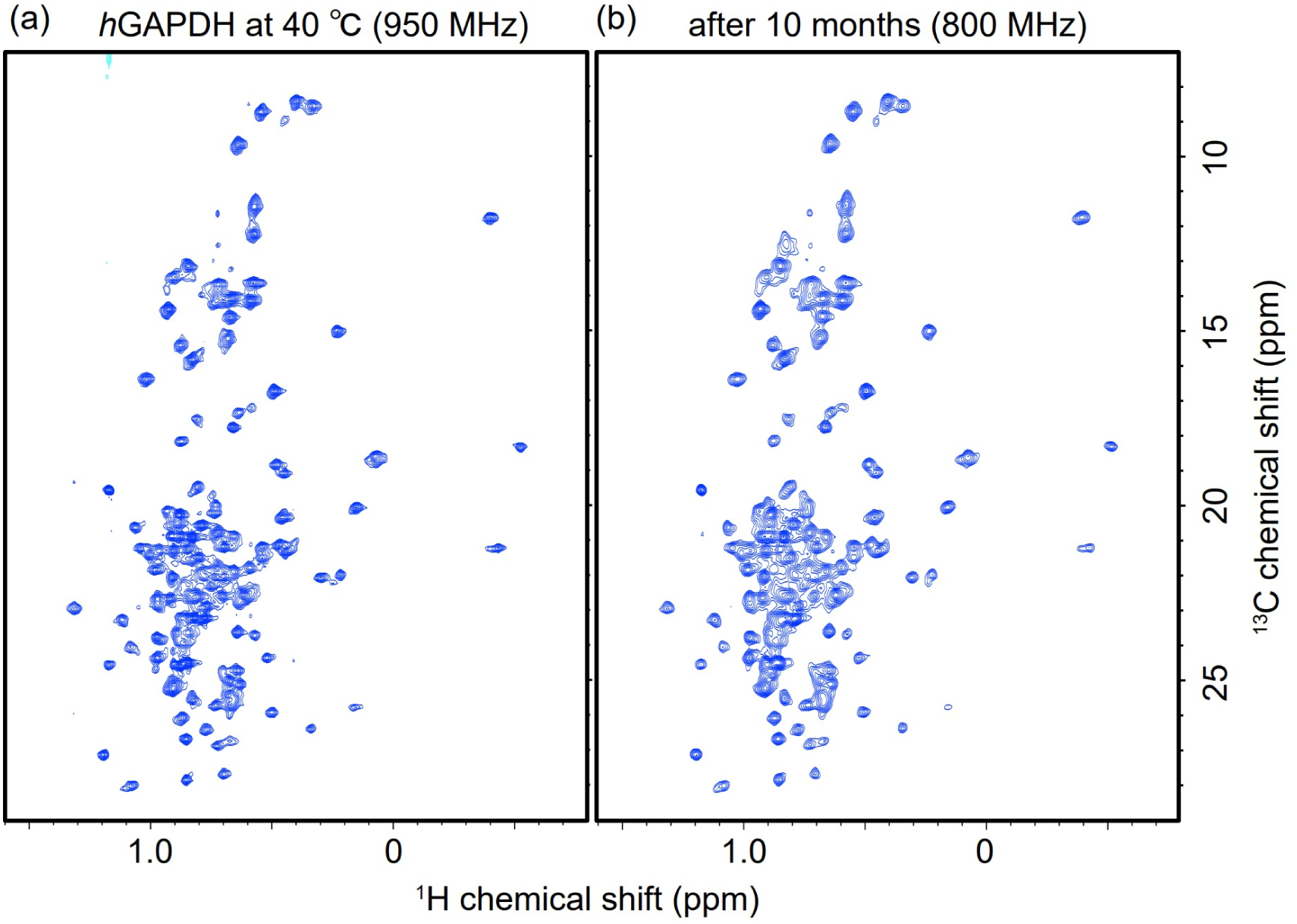
2D ^1^H-^13^C methyl-TROSY HMQC spectra of 4.0 µM {*u*-[^2^H, ^15^N]; Ile^δ^^1^-[^13^CH_3_]; Met^ε^-[^13^CH_3_]; Leu^δ^, Val^γ^-[^13^CH_3_, ^12^CD_3_]}-hGAPDH. The protein was dissolved in 20 mM sodium phosphate buffer (pH 7.4) containing 10 mM NaCl, 1 mM EDTA, 1 mM DTT, and 10% (*v*/*v*) D_2_O for locking with the addition of 2 mM NAD^+^. (a) The spectrum is identical to that shown in Figure 3a recorded at ^1^H resonance frequency of 950 MHz and temperature of 40 °C. (b) The spectrum of the same sample was obtained at 800 MHz ^1^H resonance frequency after storage at 4 °C for 10 months. The peaks in (b) appear slightly broader than those in (a), probably due to the lower resolution of the 800 MHz spectrometer compared with that of the 950 MHz spectrometer. However, almost no degradation of *holo* hGAPDH was observed over time.

We also conducted a thermal shift assay to compare the thermal stability of hGAPDH with and without NAD^+^. The assay revealed that *holo* hGAPDH started to denature at approximately 65 °C (Figure 9a, dark red), whereas the *apo* form began at approximately 50 °C (black), resulting in melting temperatures (*T*_m_) of 68.5 and 58.0 °C (Table 1). This confirmed the extreme stability of the *holo* form, in which NAD^+^ molecules bind to all four subunits, as previously reported ^25^. The melting curve of untreated hGAPDH, without any pretreatment such as the addition or removal of NAD^+^, exhibited a *T*_m_ value of approximately 61 °C, which falls between those of the *apo* and *holo* forms, with a variation of approximately ±2 °C depending on the sample lot. This variability may have arisen because of differences in the oxidation state or the degree of NAD^+^ binding saturation in the prepared hGAPDH samples.

**Figure 9:**
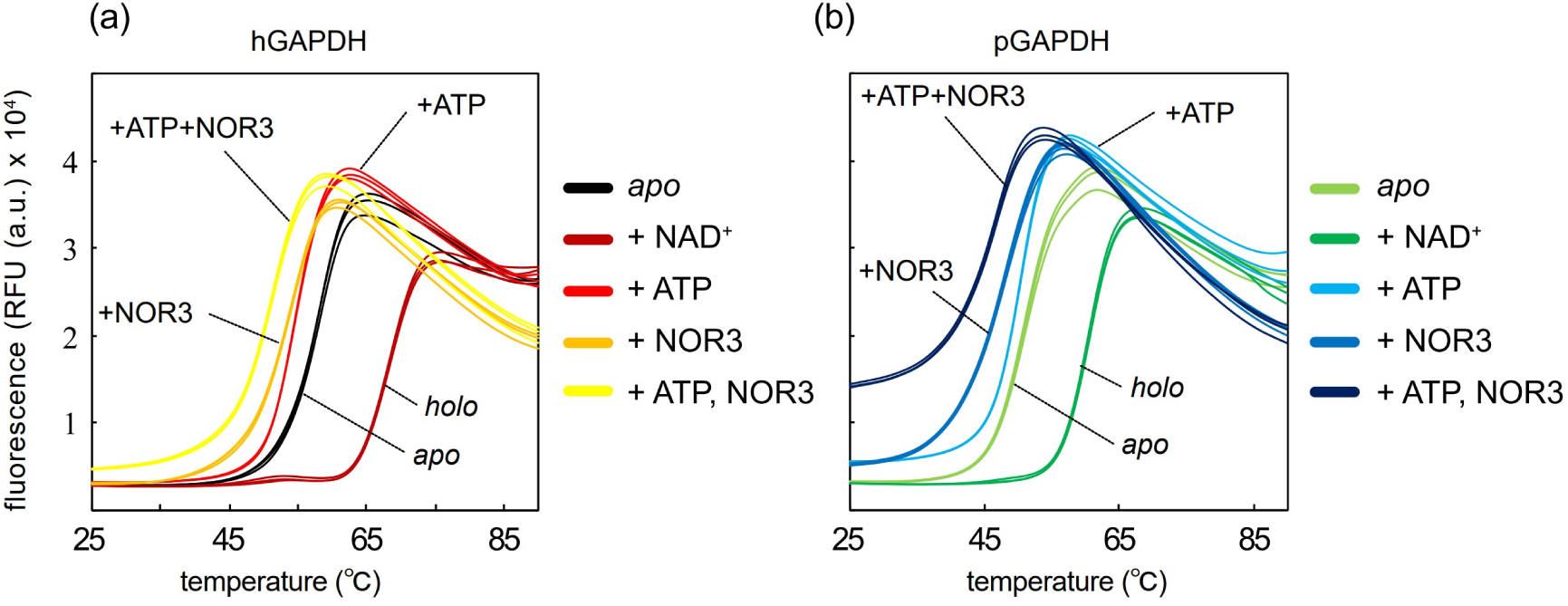
Thermal shift assays of hGAPDH (a) and pGAPDH (b) (10 µM each). The curves represent *apo* hGAPDH (black), *holo* hGAPDH with 1 mM NAD^+^ (dark red), *apo* hGAPDH with 1 mM ATP (red), *apo* hGAPDH with 1 mM (±)-(E)-4-ethyl-2-[(E)-hydroxyimino]-5-nitro-3-hexenamide (NOR3; orange), and *apo* hGAPDH with 1 mM ATP and 1 mM NOR3 (yellow) for (a) and *apo* pGAPDH (light green), *holo* pGAPDH with 1 mM NAD^+^ (green), *apo* pGAPDH with 1 mM ATP (cyan), *apo* pGAPDH with 1 mM NOR3 (blue), and *apo* pGAPDH with 1 mM ATP and 1 mM NOR3 (deep blue) for (b).

**Table 1:** Melting temperatures (*T*_m_) estimated via thermal shift assays of human and porcine glyceraldehyde-3-phosphate dehydrogenases (GAPDHs).

| hGAPDH | $T_m$ (°C) | $\Delta T_m$ (°C)<sup>a</sup> | pGAPDH | $T_m$ (°C) | $\Delta T_m$ (°C)<sup>a</sup> |
| --- | --- | --- | --- | --- | --- |
| –NAD<sup>+</sup> (*apo*)<sup>b</sup> | 58.0 | 0 | –NAD<sup>+</sup> (*apo*)<sup>b</sup> | 51.0 | 0 |
| +NAD<sup>+</sup> (*holo*)<sup>b</sup> | 68.5 | 10.5 | +NAD<sup>+</sup> (*holo*)<sup>b</sup> | 60.5 | 9.5 |
| +ATP<sup>b</sup> | 54.5 | –3.5 | +ATP<sup>b</sup> | 50.5 | –0.5 |
| +NOR3<sup>b</sup> | 54.0 | –4.0 | +NOR3<sup>b</sup> | 48.3 | –2.7 |
| +ATP+NOR3<sup>b</sup> | 51.3 | –6.7 | +ATP+NOR3<sup>b</sup> | 46.8 | –4.2 |

<sup>a</sup> Difference in each $T_m$ value from that of its corresponding *apo* form is indicated.
<sup>b</sup> Concentration of each additive is indicated in the Figure 9 legend.

### pGAPDH was less stable than hGAPDH, irrespective of the *apo* or *holo* form

We compared the stability of pGAPDHs and hGAPDHs in both the *apo* and *holo* forms using thermal shift assays. As illustrated in Figure 9 and Table 1, both the human and porcine *holo*-types exhibited approximately 10 degrees higher thermal stability than their respective *apo*-types. Interestingly, both the *apo*- and *holo*-types of pGAPDH were approximately 7–8 degrees less stable than the corresponding types of hGAPDH. This finding explains the higher number of reports demonstrating significant structural changes, including subunit dissociation, resulting from environmental factors in rabbit GAPDH and pGAPDH than those in hGAPDH.

### S-nitrosocysteine (CysNO) nitrosylated approximately half of the hGAPDH subunits

It has been reported that nitrosylation of the active site cysteine leads to the interaction of GAPDH with Siah1 and its translocation to the nucleus ^7^. Initially, we assessed the efficiency of the nitrosylation reaction by ESI-MS spectrometry under denaturing conditions to detect an increase in molecular mass by 29 u, corresponding to the covalent binding of a nitric oxide group (-NO) to the thiol group in a Cys residue. The CysNO-treated sample displayed a peak corresponding to a molecular mass 29 u higher than that of monomeric unmodified hGAPDH, whereas the untreated sample showed no such peak (Figure 10). Although CysNO demonstrated the potential for nitrosylation, a peak corresponding to the original molecular mass of unmodified hGAPDH was also observed at approximately the same intensity. This indicates that, despite employing 50-fold equivalents of CysNO, only approximately half of the hGAPDH molecules were nitrosylated.

**Figure 10:**
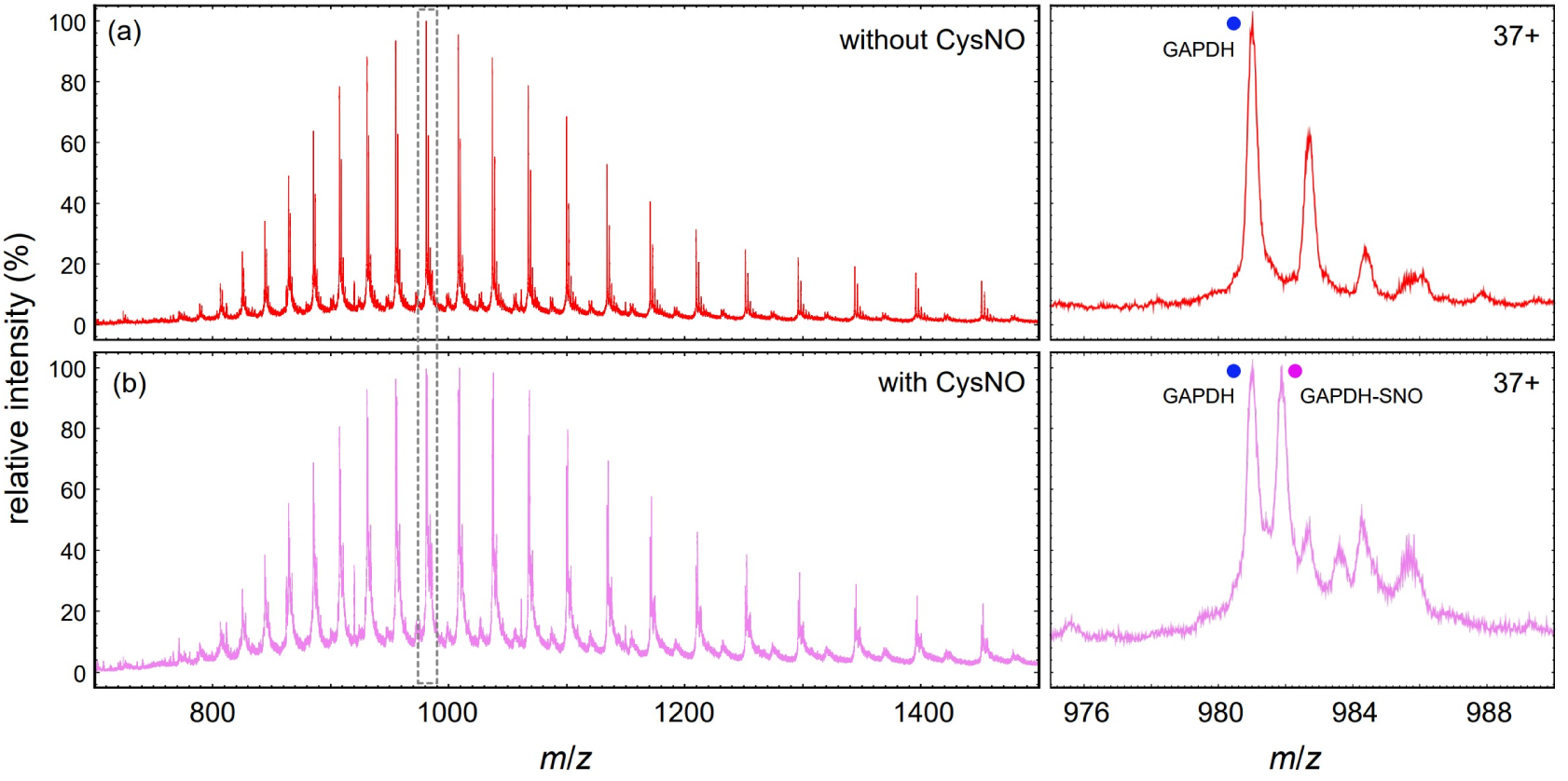
ESI mass spectra of hGAPDH before (a) and after (b) treatment with CysNO. Prior to measurement, hGAPDH was denatured in an aqueous solution containing 0.1% formic acid and 50% acetonitrile. Enlarged views of the peak regions corresponding to the 37+ charge state, outlined by dotted lines, are shown to the right of each panel. Peaks with blue circles correspond to the unmodified hGAPDH monomer, while those with magenta circles correspond to the mono-nitrosylated monomer.

### Nitrosylation of the active site Cys residue reduced NAD^+^ affinity

The ESI-MS spectra of hGAPDH, with and without CysNO treatment under native-like conditions, displayed multiple broadened peaks. The mass difference between each pair of neighboring peaks was consistent with NAD^+^ (663.4), indicating that these peaks correspond to hGAPDH with varying numbers of bound NAD^+^ molecules (Figure 2c). Non-nitrosylated hGAPDH tetramer was found to bind 1–4 NAD^+^ molecules (primarily 4 NAD^+^ molecules), whereas the nitrosylated form was found to bind 0–3 NAD^+^ molecules (primarily 2 NAD^+^ molecules). This suggests that nitrosylation reduced the affinity of GAPDH for NAD^+^. Importantly, the spectra exclusively exhibited peaks corresponding to homotetrameric hGAPDH. Furthermore, gel filtration chromatography at 25 °C revealed that nitrosylation did not alter the elution position (Figure 11). These findings indicate that mild nitrosylation, specifically targeting only the active site cysteine residue, does not cause significant conformational changes, such as subunit dissociation.

**Figure 11:**
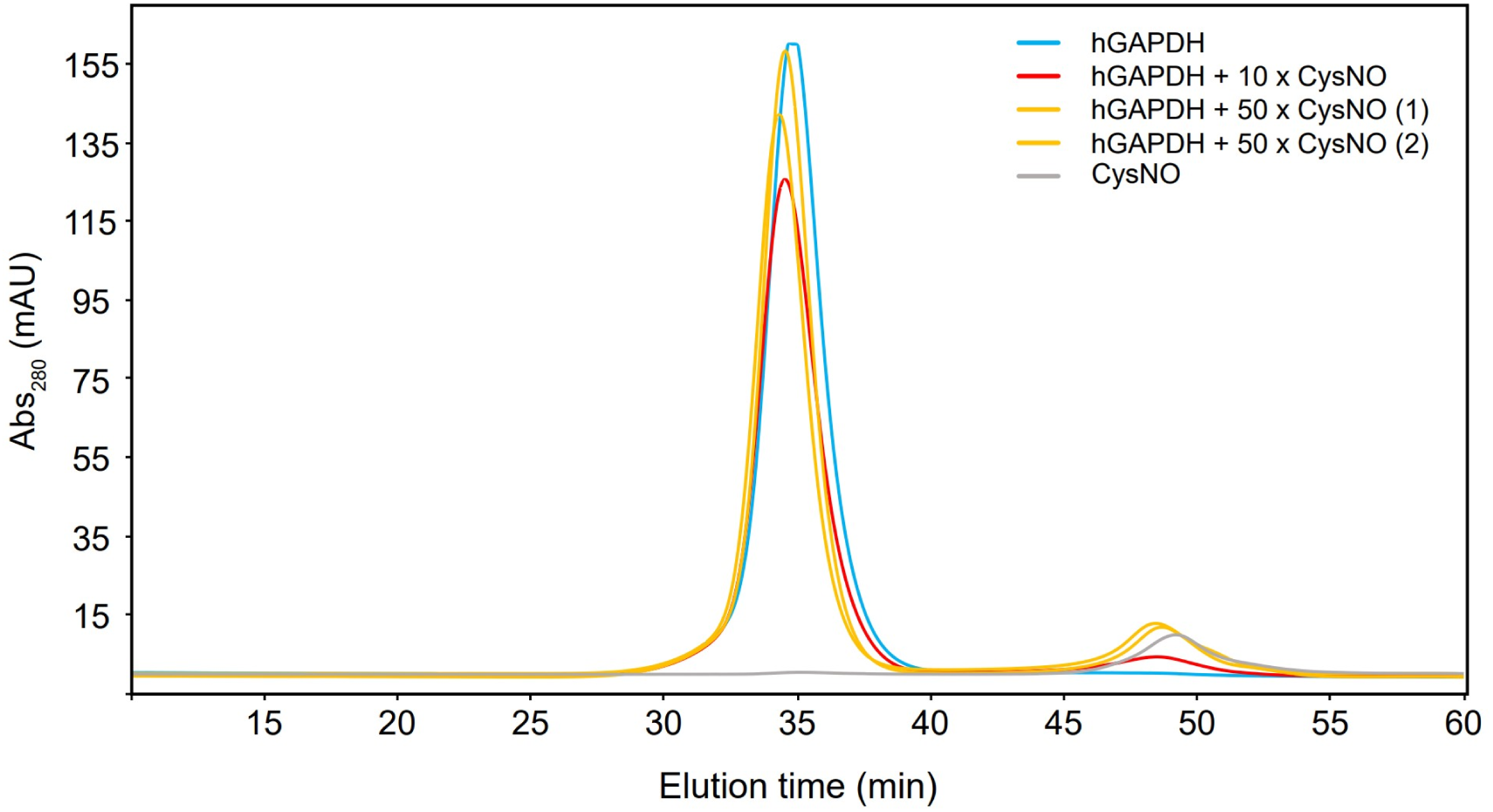
Analytical gel filtration of nitrosylated hGAPDH. *Apo* hGAPDH (cyan) and *apo* hGAPDH nitrosylated with a 10-fold equivalent (red) and 50-fold equivalent (yellow) of CysNO and with CysNO alone (grey) were individually applied to an analytical gel filtration column (Superdex 200 10/300GL) 25 °C at a flow rate of 0.4 mL/min.

### CysNO treatment caused no major conformational changes in hGAPDH

2D ^1^H-^13^C HSQC experiments on [^15^N, ^13^C]-hGAPDH nitrosylated with CysNO revealed a spectrum similar to that of unmodified *apo* hGAPDH (Figure 12). MS analysis showed that the reaction with CysNO was weak, with only half of the subunits being nitrosylated. Therefore, local nitrosylation, presumably of the cysteine residue at the active site, did not cause major conformational changes, such as subunit dissociation, at least under the NMR measurement conditions.

**Figure 12:**
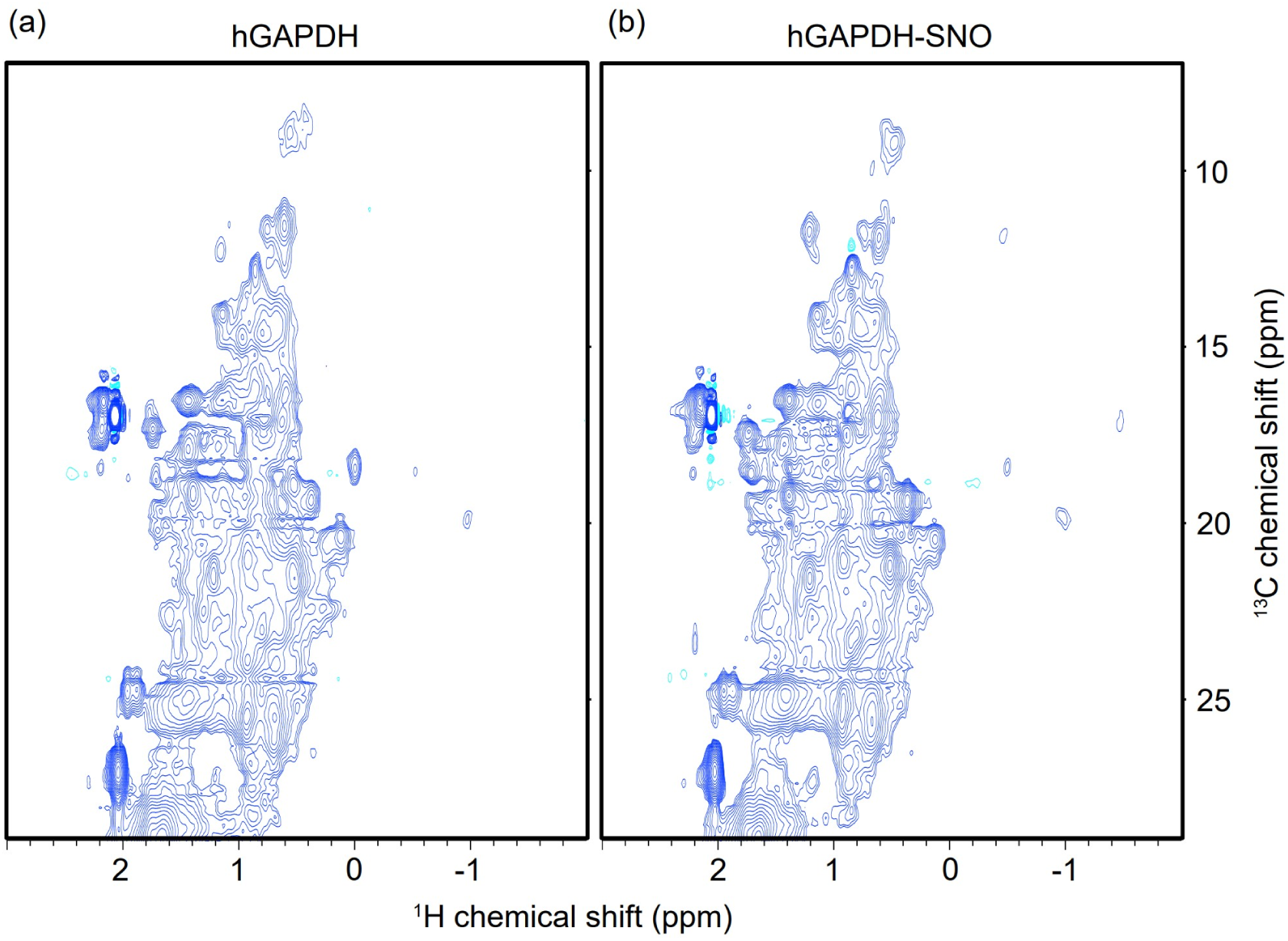
2D ^1^H-^13^C HSQC spectra of (a) *apo* [^15^N, ^13^C]-hGAPDH treated with activated charcoal and (b) *apo* [^15^N, ^13^C]-hGAPDH treated also with the 50-fold equivalent of CysNO for 30 min. For each sample, the solvent was exchanged with 20 mM Tris-HCl buffer (pH 8.0) containing 100 mM NaCl and 10% D_2_O to obtain a subunit concentration of 150 µM. The spectra were then measured at 293 K using a 500 MHz spectrometer.

### Oxidation induced precipitation and aggregation in hGAPDH

We investigated the influence of a reducing agent on the conformation of the *apo* state of hGAPDH after removing NAD^+^ using activated charcoal. Notably, the NMR experiments depicted in Figures 3, 5, and 8 used samples containing 1 mM dithiothreitol (DTT). Without DTT, we observed a significant reduction in the NMR peak intensity of the *apo* state, showing an average decrease of 65% at the beginning of the measurement at 40 °C compared to samples containing 20 mM DTT (Figure 13). This decline in peak intensity can be attributed to non-specific oxidation-induced structural changes in hGAPDH, resulting in the formation of disulfide bonds within ^26^ and between subunits, ultimately leading to aggregation and precipitation, which are typically invisible in solution NMR spectroscopy because of their high molecular weights. Furthermore, we noted that storing the measured hGAPDH sample in an NMR tube at 4 °C for one week mitigated the decrease in peak intensity to 52%, likely due to the low temperature and decreased oxygen influx.

**Figure 13:**
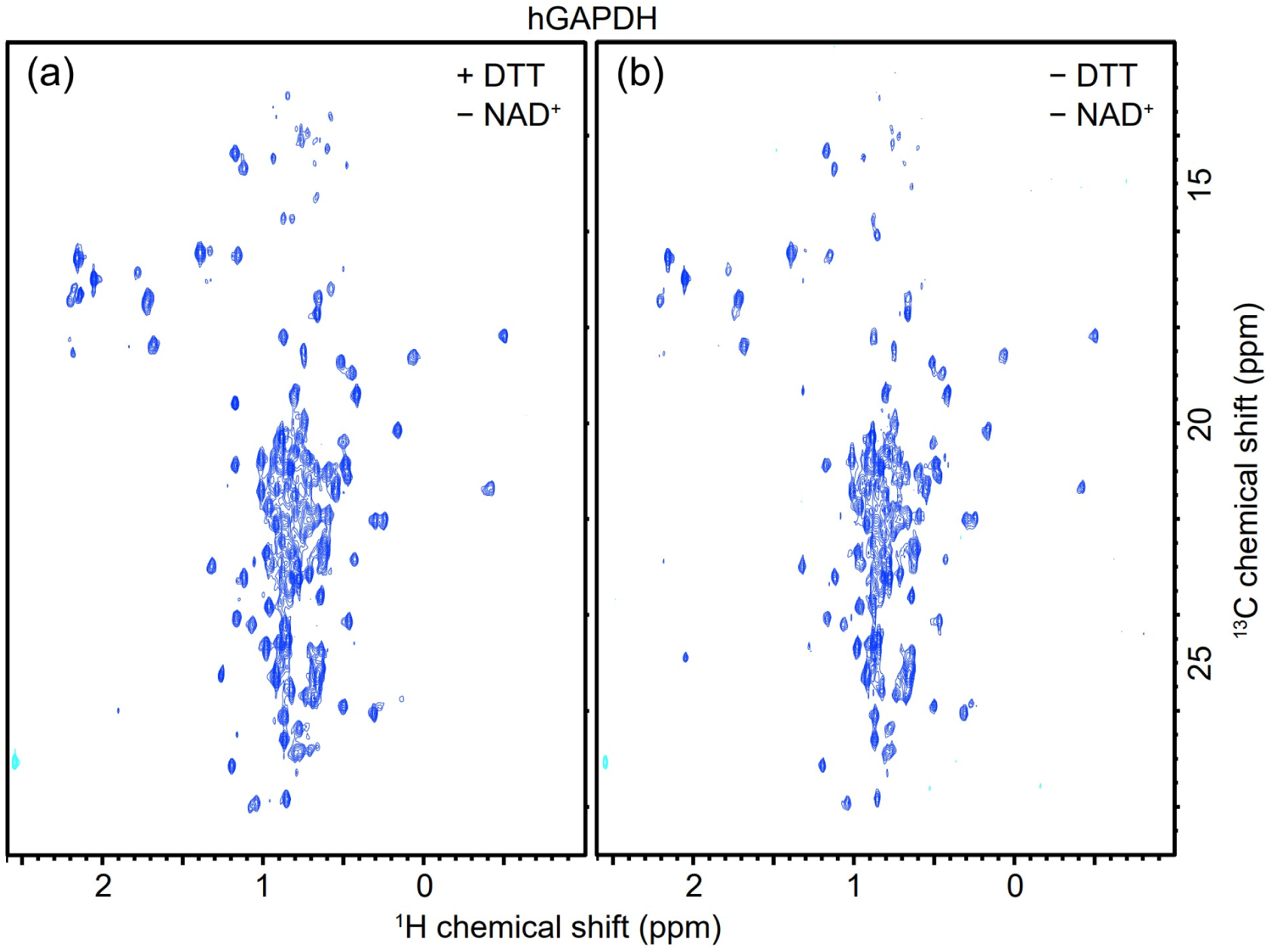
Methyl-TROSY HMQC spectra of 4.0 µM {*u*-[^2^H, ^15^N]; Met-[^13^C, ^1^H]; Leu^δ^, Val^γ^-[^13^CH_3_, ^12^CD_3_]}-*apo* hGAPDH in the presence (a) and absence (b) of 20 mM deuterated DTT. Both samples were devoid of NAD^+^. Spectra were acquired using a 950 MHz NMR spectrometer at 40 °C.

### Nitrosylation by NOR3 treatment reduced thermal stability

We used (±)-(E)-4-ethyl-2-[(E)-hydroxyimino]-5-nitro-3-hexenamide (NOR3), which exhibits a higher nitrosylating potency than CysNO, to nitrosylate *apo* hGAPDH. Figure 9a illustrates the results of the thermal shift assay conducted on two different samples: hGAPDH treated with activated charcoal alone (black) and hGAPDH treated with both activated charcoal and NOR3 (orange). The obtained results indicated that the *T*_m_ value for the *apo* hGAPDH sample without NOR3 treatment was 58 °C, whereas it decreased to 54 °C upon nitrosylation with NOR3. This observation suggests that nitrosylation of hGAPDH via NOR3 treatment leads to reduced thermostability.

### Oxidation of various residues by NOR3 caused significant structural changes in hGAPDH

Next, we nitrosylated hGAPDH with NOR3, which exhibits better nitrosylation and oxidation activities than CysNO, to investigate the impact of NOR3 on the protein structure. Figure 14 shows the ^1^H-^13^C methyl-TROSY HMQC spectra of *apo* hGAPDH with and without NOR3 treatment. The sample without NOR3 displays distinct, rounded, and sharp NMR peaks. In contrast, the sample treated with NOR3 exhibits broadened peaks, with some peaks even disappearing. Typically, such degradation of NMR peaks may arise because of denaturation, aggregation, increased flexibility (on a slow and intermediate NMR timescale), or dynamic exchange between multiple intra- and intermolecular structures. NOR3 is recognized for its ability to oxidize several residues of GAPDH, including Tyr and Trp, and nitrosylate the cysteine residue at its active site ^27^. Our NMR results suggest that these oxidative modifications induce significant structural alterations in *apo* hGAPDH.

**Figure 14:**
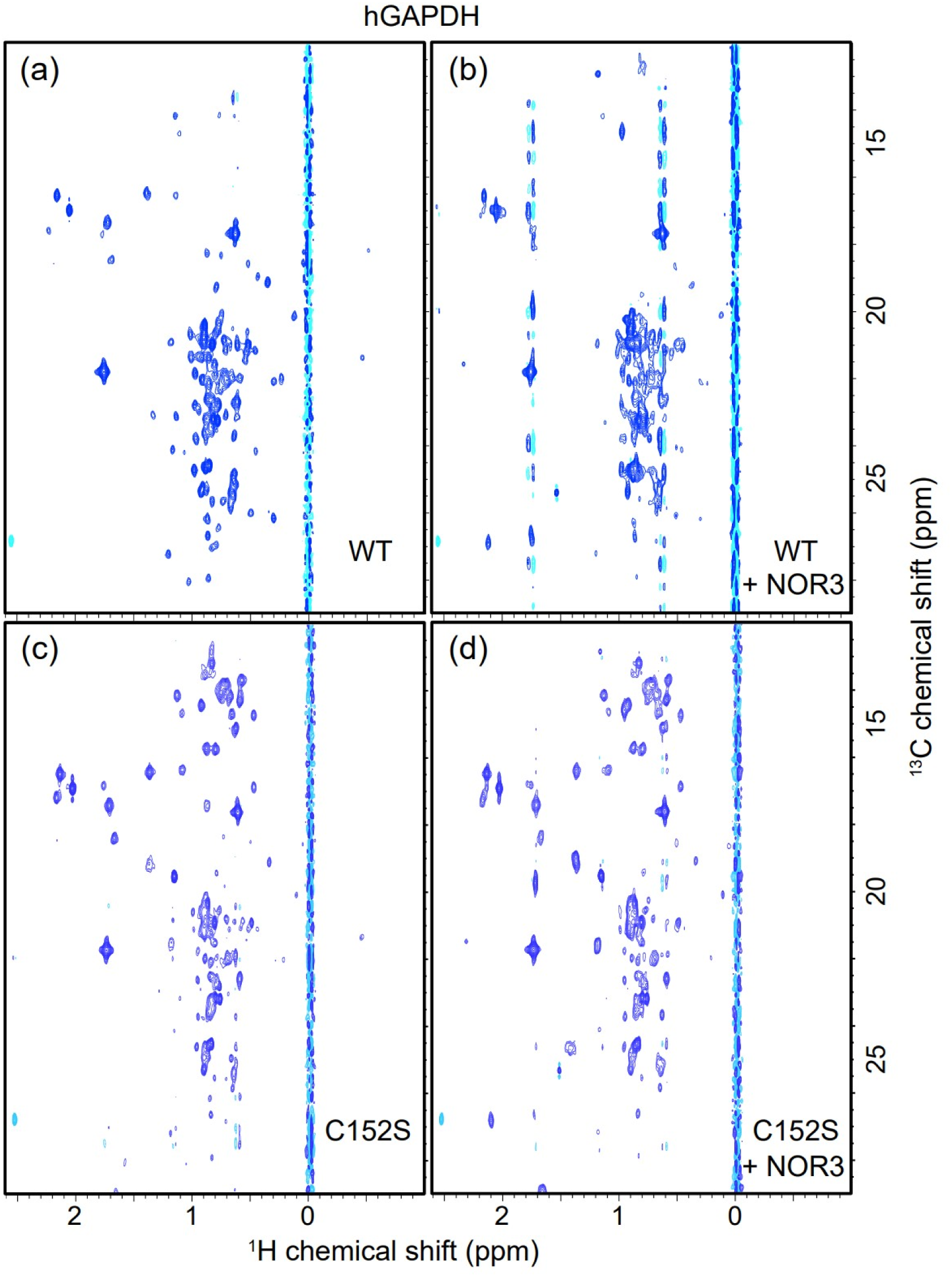
Methyl-TROSY HMQC spectra of 4.0 µM {*u*-[^2^H, ^15^N]; Ile^δ^^1^-[^13^CH_3_]; Met^ε^-[^13^C]; Leu^δ^, Val^γ^-[^13^CH_3_, ^12^CD_3_]}-*apo* hGAPDH wild-type (wt) (a, b) and C152S variant (c, d) in the absence (a, c) and presence (b, d) of 100 µM NOR3. The reference samples (a, c) contained the same amount of stock solvent (1% deuterated dimethyl sulfoxide [*d*-DMSO]) as those in (b, d). The solvent was 20 mM sodium phosphate buffer (pH 7.4) containing 10 mM NaCl, 1 mM EDTA, 1 mM DSS, and 10% D_2_O. The spectra were measured using an 800 MHz NMR spectrometer at 25 °C. In the preparation of the wt samples, the amount of Ile precursor, [3,3-^2^H_2_, 4-^13^C_1_]-α-ketobutyrate, added to the minimal medium for *E. coli* culture was lower than required (130 mg/L instead of 200 mg/L), resulting in weak visibility of the δ-methyl peaks of Ile.

### ATP destabilized the tetrameric conformation of both porcine and human GAPDH

NAD^+^ removal and local nitrosylation of GAPDH alone did not cause significant conformational changes according to our NMR measurements at room temperature. However, thermal shift assays clearly demonstrated that these events reduced stability. This may be because a small molar ratio of GAPDH underwent conformational changes or subunit splitting. Such low-abundance structures are generally difficult to detect using simple NMR measurements, which require an increase in population. Previous studies have reported that adding ATP to rabbit, yeast, pig, and *E. coli* GAPDHs promotes subunit dissociation into dimers and monomers, indicating that ATP destabilizes the tetramer conformation. ^28–32^. Therefore, we conducted conformational analysis of GAPDH in the presence of ATP, anticipating that ATP would increase the molar ratio of conformationally altered GAPDH. The results of the thermal shift assays for hGAPDH and pGAPDH are presented in Figure 9, and the obtained *T*_m_ values are listed in Table 1. ATP significantly decreased the thermal stability of both hGAPDH and pGAPDH, with denaturation temperatures 3.5°C and 0.5°C lower than their respective *apo* forms.

### Addition of ATP to *apo* pGAPDH at low temperatures led to subunit dissociation

Analytical gel filtration was used to determine whether ATP or temperature affects the quaternary structure of GAPDH. Figure 15a shows the elution profile of *holo* pGAPDH in the presence of NAD^+^. Another analytical gel filtration (Figure 6) revealed that pGAPDH eluted as a tetramer, regardless of the presence or absence of NAD^+^ at both 4 °C and 28 °C. However, when ATP was added to *apo* pGAPDH, it eluted at a lower molecular weight at 4 °C (Figure 15b), but as a tetramer at 28 °C (Figure 15c). A closer examination of the elution profiles revealed that, even at 28 °C, a small peak was observed at a lower molecular weight, and at 4 °C, a portion of the elution was observed at the tetramer position. These results suggest that the tetrameric structure becomes less stable in the *apo* state and that this instability is exacerbated at lower temperatures. Furthermore, ATP stabilizes dimers or monomers ^31^, shifting the pGAPDH equilibrium towards subunit dissociation owing to low-temperature denaturation.

**Figure 15:**
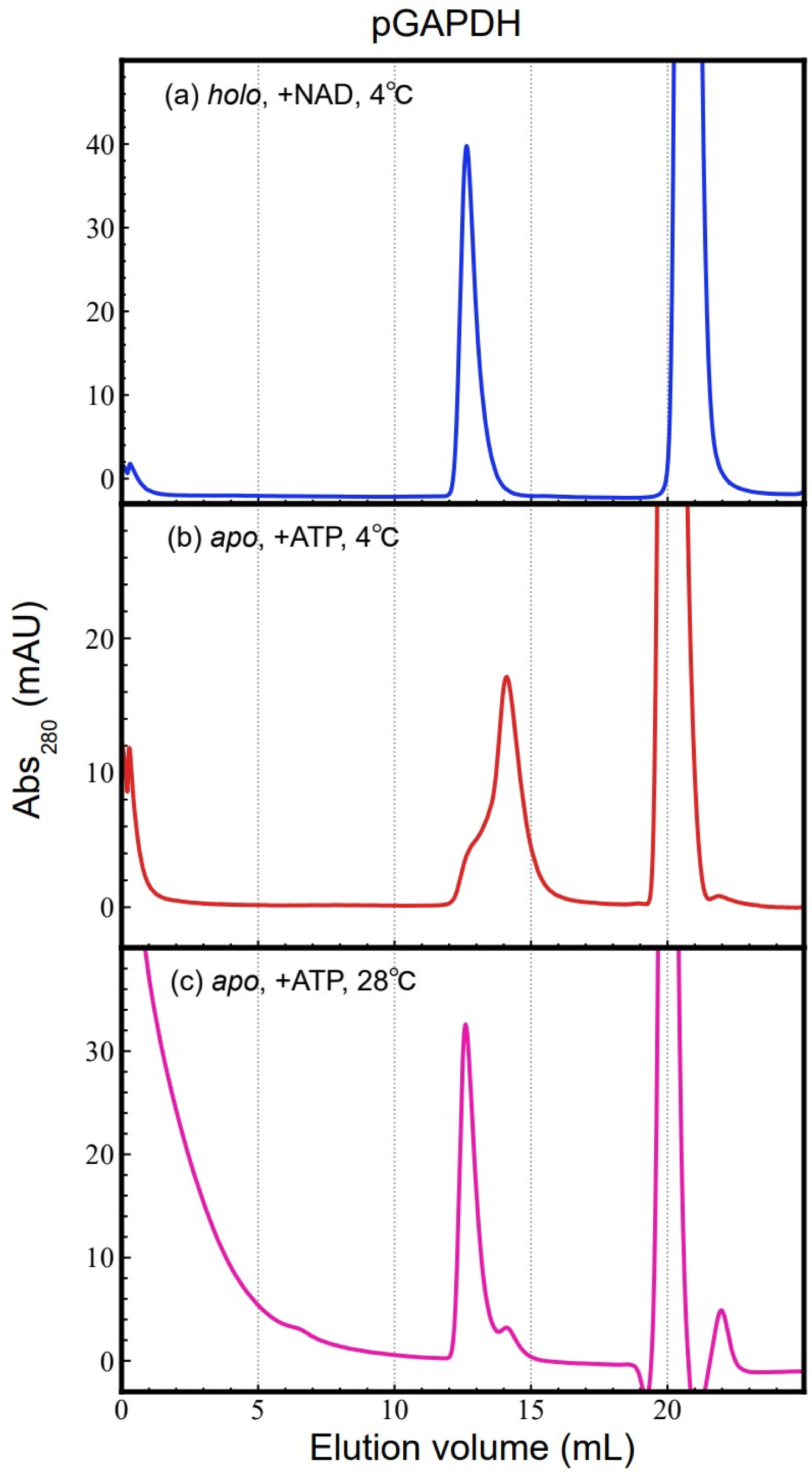
Analytical gel filtration of pGAPDH. pGAPDHs (25 µM, 300 µL) with (*holo*) and without (*apo*) NAD^+^ were separately applied to an analytical gel filtration column (Superdex 200 increase 10/300 GL) at 4 and 28 °C at a flow rate of 0.38 mL/min. The running buffer comprised 20 mM Tris-HCl (pH 8.0) and 200 mM NaCl. (a) To analyze *holo* pGAPDH at 4 °C, the injected sample contained 2 mM NAD^+^, and the running buffer also contained 1 mM NAD^+^. NAD^+^ was eluted in a volume of approximately 21 mL. (b) *Apo* pGAPDH was incubated with 2 mM ATP for 12 h at 4 °C and analyzed at the same temperature using a running buffer supplemented with 1 mM ATP. (c) *Apo* pGAPDH was incubated with 2 mM ATP for 12 h at 28 °C and analyzed at the same temperature using a running buffer supplemented with 1 mM ATP.

### pGAPDH exhibited flexible residues at low temperatures in the presence of ATP

We obtained a 2D ^1^H-^13^C NMR spectrum of *apo* pGAPDH with ATP at 25 °C. Comparison with the spectrum without ATP (Figure 16a,b) showed no significant differences, indicating that, at room temperature, *apo* pGAPDH retained its normal tetrameric conformation, even with ATP, which is consistent with the gel filtration analysis results (Figure 15c). However, at 5 °C, NMR spectra of the same samples differed in the presence and absence of ATP (Figure 16c,d). Although most peaks showed no significant chemical shift differences, sharper peaks were observed for the sample containing ATP. These peaks overlapped with those of free Met, Ile, Leu, and Val amino acids (Figure 16e), suggesting that ATP interactions induce partial unfolding and increase the flexibility in specific regions. Interestingly, these peaks were distorted along the ^13^C dimension, a phenomenon typically observed when the signal intensities (time-domain data along the ^13^C dimension) gradually increase during measurement. Therefore, more molecules developed flexible regions over time (18 h). After measuring the spectrum at 5 °C, the temperature was returned to 25 °C, and measurements were repeated. As shown in Figure 16f, sharp peaks disappeared, and the spectrum returned to its pre-5 °C state (Figure 16b). These results suggest that ATP interactions at low temperatures reversibly increase the number of flexible regions in the GAPDH conformation.

**Figure 16:**
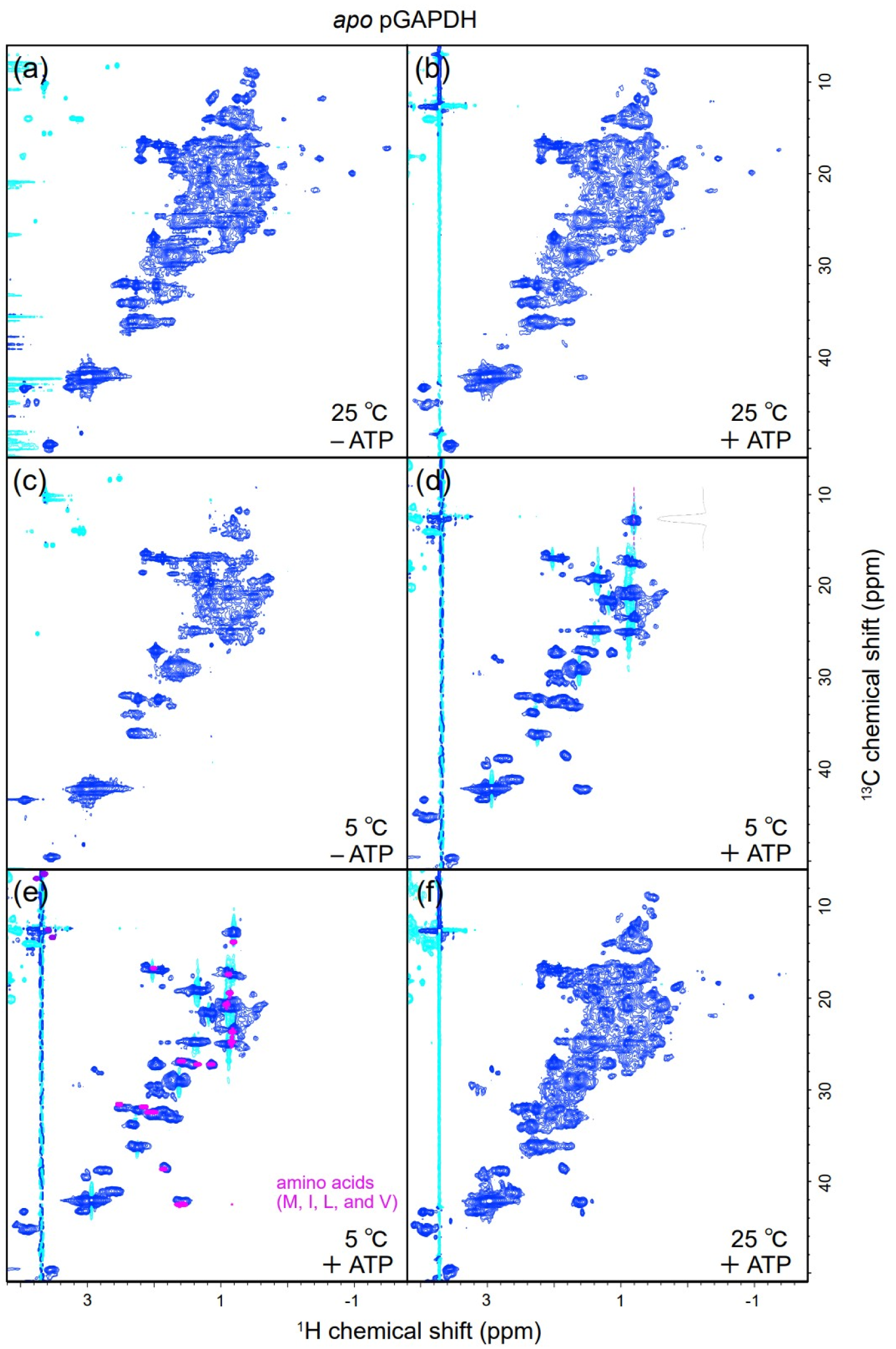
2D ^1^H-^13^C HSQC spectra of *apo* pGAPDH with and without ATP. (a) Solution comprised 100 µM [^15^N, ^13^C]-*apo* pGAPDH, 20 mM sodium phosphate (pH 7.4), 10 mM NaCl, 1 mM EDTA, 1 mM DTT, and 10% D_2_O. (b) Solution comprised 114 µM [^15^N, ^13^C]-*apo* pGAPDH, 20 mM Tris-HCl (pH 6.8), 10 mM NaCl, 10% D_2_O, and 10 mM ATP. Spectra of (a) and (b) were acquired at 25 °C. Immediately after the measurements, spectra of (c) and (d) were acquired at 5 °C. Cyan contours indicate negative intensity. An example of the distorted peak shape is shown as a one-dimensional cross-section at the magenta dotted line for the most ^13^C low-frequency peak. (e) Spectrum of (d) overlaid with that of free amino acids (magenta). For the latter, Met, Ile, Leu, and Val powders were dissolved in 100 mM HCl and 7% D_2_O, and the spectrum was measured at 30 °C. (f) A spectrum was acquired again at 25 °C after measurements at 25 °C (b) and 5 °C (d).

### pGAPDH unfolded irreversibly and synergistically in the presence of both ATP and NOR3

A similar experiment was conducted for hGAPDH; however, the NMR spectrum of *apo* hGAPDH in the presence of ATP at 5 °C did not exhibit changes as clear as those observed for pGAPDH. This finding is consistent with the thermal shift assay result that hGAPDH was more stable than pGAPDH. Nevertheless, both variants exhibited similar characteristics, with either ATP or NOR3 treatment reducing the *T*_m_ values of GAPDHs of both species (Figure 9; Table 1). Furthermore, simultaneous presence of ATP and NOR3 led to greater destabilization of hGAPDH and pGAPDH, causing a decrease in *T*_m_ values by 6.7 and 4.2 °C, respectively, compared to those of their respective *apo* forms. Therefore, combined effects of ATP-interaction and NOR3-oxidation shift the conformational equilibrium of GAPDHs towards a partially unfolded state, resulting in a synergistic destabilization effect. To examine the effect on conformation, NMR measurements of *apo* pGAPDH were conducted in the presence of ATP and NOR3. As shown in Figure 17, the spectrum obtained at 5 °C showed strong peaks derived from the unfolded regions, similar to the spectrum of *apo* pGAPDH with ATP alone (Figure 16d). Notably, after the measurement at 5 °C, these strong peaks persisted even when the temperature was increased to 25 °C (Figure 17b). In contrast, in the presence of ATP, but not NOR3, increasing the temperature to 25 °C caused the spectrum to revert to that observed in the absence of ATP (Figure 16f). These results indicate that NOR3 treatment in the presence of ATP irreversibly unfolds *apo* pGAPDH. Moreover, ATP induces a quaternary structural change from a tetramer to a dimer, which is prone to oxidation by NOR3.

**Figure 17:**
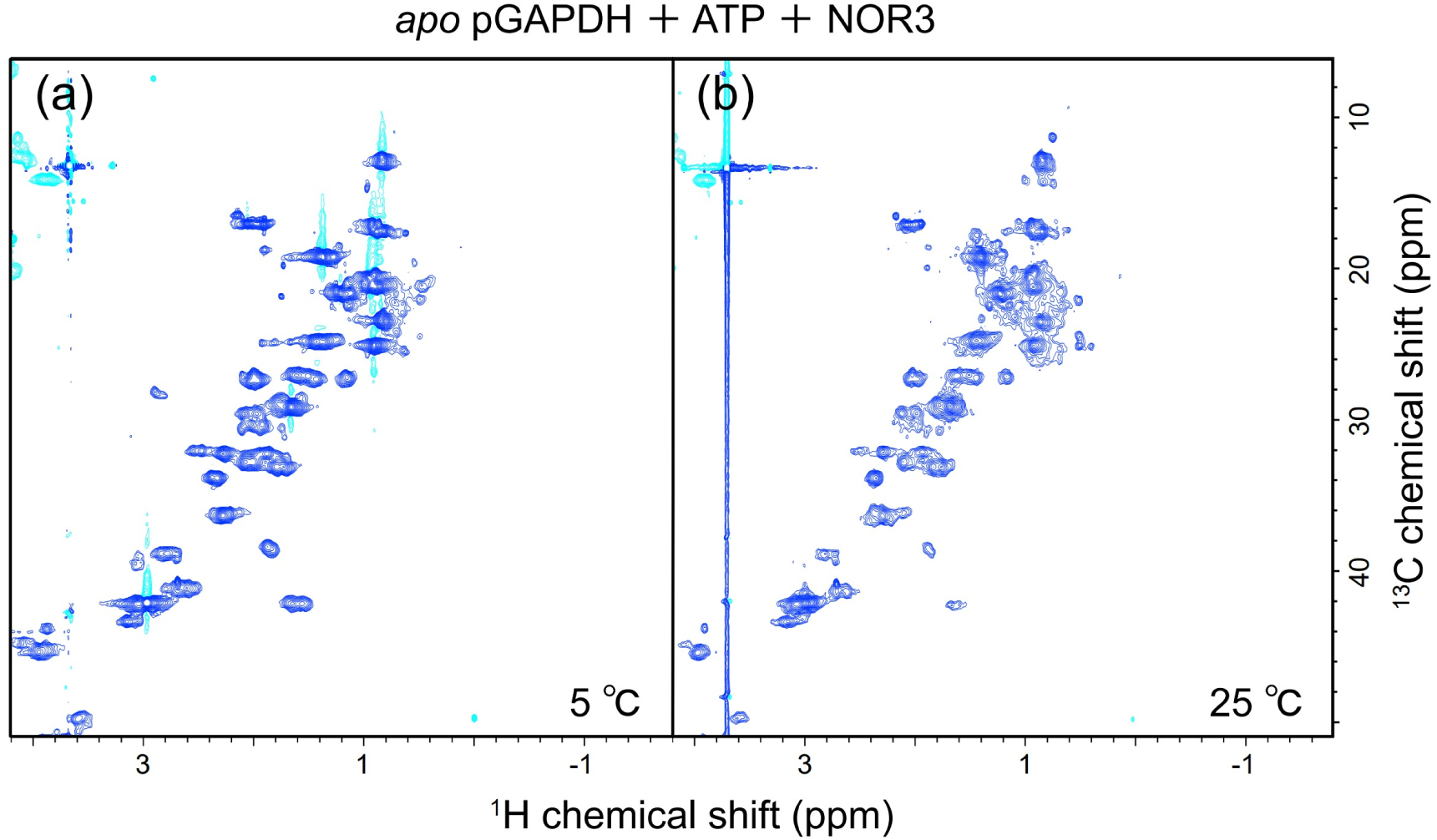
2D ^1^H-^13^C HSQC spectra of *apo* pGAPDH in the presence of ATP and NOR3. The solution comprised 77 µM [^15^N, ^13^C]-*apo* pGAPDH, 20 mM Tris-HCl (pH 8.5), 200 mM NaCl, 10 mM ATP, 100 µM NOR3, 0.1 mM DSS, and 10% D_2_O. Spectra were measured at 5 °C (a) and then at 25 °C (b).

### Preliminary NMR experiments suggested weak interactions between GAPDH and other molecules

To investigate the interactions of GAPDH with other molecules, we labeled GAPDH or its interacting partners with stable isotopes (^15^N and ^13^C) and analyzed their interactions by 2D ^1^H/^15^N or ^1^H/^13^C HSQC experiments. In these spectra, NMR peaks arise solely from the ^13^C- and ^15^N-labeled molecules, while unlabeled partner molecules remain silent. While the NMR chemical shift assignment of hGAPDH remains incomplete, we provide a preliminary overview of three ongoing experiments. Comprehensive results will be detailed in the future.

Initially, we labeled α-synuclein with ^15^N and mixed it with unlabeled *apo* hGAPDH that had been treated with CysNO to monitor any possible changes in the amide group ^1^H/^15^N chemical shifts of α-synuclein. As shown in Figure 18a, no significant chemical shift perturbations were observed under the current experimental conditions. However, we noted a substantial decrease in peak intensity (approximately to 60%) in the region from residue 121 to the C-terminus of α-synuclein. This decrease suggests a transient and weak interaction between the C-terminal region of α-synuclein and hGAPDH. Given the reported interaction between oxidized GAPDH and α-synuclein in previous studies ^20^, we are currently investigating this interaction using a similar experimental approach with GAPDH oxidized with H_2_O_2_.

**Figure 18:**
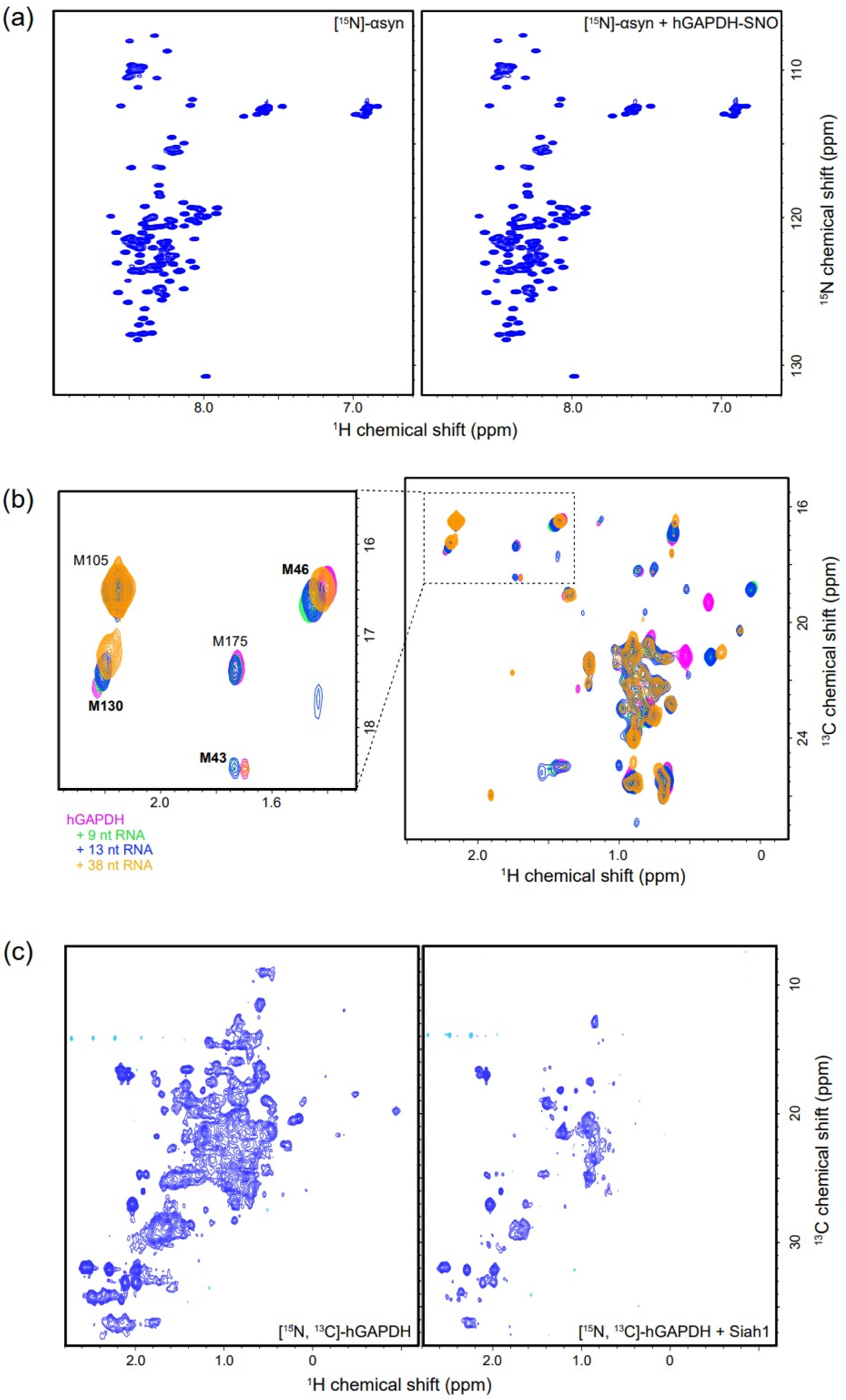
(a) 2D ^1^H-^15^N HSQC spectra of 90 μM [^13^C, ^15^N]-α-synuclein in the absence (left) and presence (right) of 130 μM hGAPDH-SNO. The solvent was phosphate-buffered saline (pH 7.4) containing 10% D_2_O, and the spectra were measured using a 500 MHz NMR spectrometer at 20 °C. (b) Overlay of 2D ^1^H-^13^C HSQC spectra of 50 μM *apo* [^13^C, ^15^N]-hGAPDH measured in the absence (pink) and presence of 200 μM 9-base (green), 142 μM 13-base (blue), and 49 μM 38-base (orange) RNAs. The solvent was 20 mM sodium phosphate buffer (pH 7.4) containing 10 mM NaCl, 1 mM EDTA, 0.1 mM DSS, 1 mM DTT, and 10% D_2_O. The spectra were recorded using an 800 MHz NMR spectrometer at 25 °C. The region displaying the Met methyl peaks is magnified, with tentatively assigned residue names indicated. (c) 2D ^1^H-^13^C HSQC spectra of 50 μM [^13^C, ^15^N]-hGAPDH, untreated with activated charcoal, in the absence (left) and presence (right) of 150 μM Siah1 (C16S/C282S variant). The solvent was 25 mM Tris-HCl buffer (pH 8.0) containing 10 mM NaCl, 1 mM DTT, 30 μM ZnCl_2_, 0.1 mM DSS, and 10% D_2_O. Spectra were recorded using an 800 MHz NMR spectrometer at 40 °C.

The mRNAs encoding cytokines frequently contain adenine-uridine rich elements (AREs), and certain RNA-binding proteins regulate the stability of target mRNAs by binding to AREs and preventing endonuclease access. Such an ARE is located in the 3’ untranslated region of tumor necrosis factor (TNF)-α mRNA, where an interaction with GAPDH has been reported ^33^. To further investigate this interaction, we mixed [^15^N, ^13^C]-hGAPDH with 38-base, 13-base, or 9-base synthetic RNAs and observed changes in the chemical shifts of several methyl groups (Figure 18b). The magnitude of these shifts was larger for longer RNAs, suggesting stronger interactions with longer RNAs. While the assignment of the methyl groups is ongoing, the shifted peaks appear to originate from the active site and positively charged groove, supporting previous reports ^34^.

Previous studies have demonstrated that S-nitrosylated GAPDH forms a complex with Siah1, leading to its nuclear translocation and the induction of apoptosis ^7^. In the present investigation, we added unlabeled, intact Siah1 to [^15^N, ^13^C]-hGAPDH and monitored changes in the ^1^H/^13^C chemical shifts of hGAPDH (Figure 18c). We hypothesized that NMR spectroscopy could detect weak interactions between these proteins, even in the absence of S-nitrosylation. Indeed, we observed shifts and broadening of methyl group peaks in hGAPDH. Future assignment of these methyl groups will enable us to identify whether the interaction is specific and, if so, where the specific interaction sites exist. AlphaFold2 ^24^ predicts that the N-terminal region (residues 1-31) of Siah1 is intrinsically disordered. On the basis of the predicted structural information, we are examining the interactions between GAPDH-SNO and Siah1 variants with varying domain configurations.

## Discussion

### NMR is well-suited for the analysis of flexible and dynamic structures

Crystal structures of GAPDHs derived from various species have been reported. However, the structure of GAPDH, when forming complexes with other proteins or nucleic acids to function as a moonlighting protein in roles beyond glycolysis, remains unclear. In many cases, cellular senescence and oxidation trigger the non-glycolytic functions of GAPDH. Under these conditions, cells have more oxidizing agents and less NAD^+^ levels than normal. Therefore, removal of NAD^+^, accompanied by further oxidation, including chemical modifications, such as glutathionylation ^26,35–38^, possibly causes conformational changes in GAPDH, allowing it to interact with other molecules. Indeed, GAPDHs in specific states, including the nitrosylated state, interact with biomolecules, such as Siah1 ^7^, amyloid-β ^39^, α-synuclein ^20^, p300/CREB binding protein ^40^, lactate dehydrogenase (LDH) ^41,42^, malate dehydrogenase 1 (MDH1) ^43^, mucin ^44^, and AU-rich elements of mRNAs ^45,46^. However, conformational changes at the atomic resolution level remain unknown. The slow progress in the structural analysis of moonlighting GAPDHs may be due to the transient or dynamic nature of their complexes. To capture their structural information, which is generally difficult to obtain via crystallographic analysis, we used NMR spectroscopy in this study.

### Methyl TROSY, in combination with deuteration, facilitates the observation of signals from high-molecular-weight proteins

NMR signals, particularly those from amide ^1^H/^15^N nuclei, are difficult to observe in high-molecular-weight proteins, such as GAPDH, even when the non-labile hydrogens are deuterated. Additionally, lower measurement temperatures slow the rotational diffusion of these molecules, further broadening the peak width and reducing their height. However, by exclusively labeling the methyl groups of Ile^δ1^, Leu, Val, and Met residues with ^1^H/^13^C and all other groups with ^2^H/^12^C, and using the methyl-TROSY mechanism ^47^, we successfully observed methyl-derived signals even at subunit concentrations as low as 1 µM and temperatures as low as 5 °C. Although assigning methyl group signals remains challenging, we are exploring this by individually mutating the methyl-containing residues. Further progress in this context will facilitate precise analysis of the conformational changes in GAPDH and its interactions with other molecules.

### NAD^+^ removal alone does not directly cause subunit dissociation, but the resulting *apo* form is more susceptible to oxidation

Cellular senescence decreases the amount of NAD^+^ ^48^ and increases the levels of reactive oxygen species (ROS) and NO in cells; artificially removing NAD^+^ from GAPDH and treating it with oxidation agents such as NOR3 *in vitro* can mimic the situation of GAPDH at the time of aging. Because GAPDH exhibits negative cooperativity in its affinity for NAD^+^, the cofactor does not easily dissociate from the homotetramer, even under conditions where the intracellular NAD^+^ level drops considerably (four dissociation constants (*K*_d_) in the range of 10^-4^-10^-9^ M ^3^). However, when cells senesce to a point where NAD^+^ is depleted, the population of the fully *apo* form increases. Our NMR and analytical gel filtration experiments showed that even when all NAD^+^ cofactors were dissociated, the GAPDH homotetramer did not immediately cleave into monomers or dimers (Figures 6 and 7). In the absence of an oxidizing agent in solution or in the presence of a reducing agent such as DTT, the *apo* form maintains a homotetrameric conformation. This indicates that NAD^+^ dissociation is not the direct cause of the large quaternary conformational change in GAPDH. Nevertheless, the thermal shift assay revealed that the *apo* form was unstable and susceptible to attack by ROS and other oxidants (Figure 9). Subsequent oxidation of multiple residues may trigger transformation into a structure associated with moonlighting functions. Furthermore, replacing NAD^+^ with ATP may shift equilibrium from tetramers to dimers or monomers. Alternatively, NAD^+^ removal and oxidation may act cooperatively, as we have shown that nitrosylation of the active site cysteine residue promotes NAD^+^ removal (Figure 2a,c).

### Although pGAPDH is less stable than hGAPDH, *apo* pGAPDH still maintains its tetrameric conformation

Ultracentrifugal analyses previously demonstrated that homotetramers dissociated into dimeric and monomeric subunits by simply removing NAD^+^ at low temperatures ^15,16,29,31^. However, our NMR and gel filtration analyses of pGAPDH and hGAPDH at low temperatures revealed that the removal of NAD^+^ alone did not dissociate the homotetramer into dimers or monomers, as long as they were kept in a reduced state in the absence of ATP. These results suggest that subunit dissociation requires further oxidation of the destabilized *apo* form or other post-translational modifications.

Many previous studies used porcine- or rabbit-derived GAPDHs extracted directly from skeletal muscle rather than those expressed in recombinant *E. coli* cells. During muscle purification processes, steps such as ammonium sulfate precipitation and crystallization followed by re-dissolution may have promoted the oxidation of GAPDHs, thereby destabilizing the proteins and facilitating the dissociation of the subunits. Therefore, we expressed GAPDHs in *E. coli* and prepared samples in a reduced state by constantly adding DTT during purification to compare their characteristics with those previously reported. Another reason for the frequent observation of subunit splitting could be that GAPDHs from pigs and rabbits are less stable than hGAPDH. Our NMR analyses of pGAPDH showed more pronounced conformational changes than those observed for hGAPDH at low temperatures in the presence of NOR3 or ATP (Figure 17). This was supported by our thermal shift assays, which revealed that pGAPDH was more unstable than hGAPDH under all the examined conditions (Table 1).

However, we must also consider that we performed almost all experiments at GAPDH concentrations above 1 μM. Lakatos *et al*. estimated the tetramer–dimer and dimer–monomer *K*_d_ values as roughly 2.2 and 1.1 µM at 5 °C, respectively, based on the concentration-dependent subunit composition of pGAPDH ^15^. A similar phenomenon was reported for GAPDH isolated from rabbits. The tetramer reversibly dissociated into dimers at 5 °C (*K*_d_ = 0.5 µM) with a subunit concentration < 14 µM (0.5 g/L) ^17^. Considering the reported equilibrium constants, it is likely that, at <1 μM and 4 °C, the *apo* form is predominantly in the form of dimers and monomers. Indeed, in our gel filtration experiments at 4 °C using *apo* pGAPDH, a small but significant elution peak was observed at the dimer position (marked with an asterisk in Figure 6).

The question that arises is whether these artificial conditions reflect those in actual mammalian bodies. GAPDH is abundant in cells, and the body temperature is maintained at 36-39°C except during hibernation. In addition, the excluded-volume effect in the crowded environment of cells tends to shift the equilibrium toward the tetramer ^49^. Conversely, the crowded environment may destabilize the GAPDH tetramer through increased collision frequency with other molecules. Furthermore, ATP-binding and post-translational modifications are likely to shift the equilibrium toward subunit dissociation. Taken together, the behavior of GAPDHs in cells may be quite different from that in purified and reduced *in vitro* environments, which may explain why the moonlighting functions and interactions with other molecules have not been observed much *in vitro*. To this end, we are analyzing the structures of GAPDH under conditions that mimic the intracellular environment.

### Inter-subunit regions may interact with other molecules

ATP functions as a molecular chaperone stabilizing *E. coli* GAPDH in its intermediate folding state. Namely, ATP shifts the structural equilibrium of GAPDH from a folded homotetrameric state to a slightly unfolded one^28^. Here, NMR and analytical gel filtration were used to examine the effect of ATP on enzyme conformation. In the presence of ATP, *apo* pGAPDH exhibited sharp and intense peaks at low temperatures. These peaks were located close to the chemical shift values of free amino acid residues (Figure 16e). When GAPDH split into dimers or monomers, the side chains in the interfacial regions, which are packed in the homotetrameric state, became solvent-exposed and flexible. The NMR peaks of such flexible side chains often appear similar to those of free amino acids. In our analytical gel filtration in the presence of ATP at low temperatures, a fraction of *apo* GAPDH was eluted at a molecular weight equivalent to that of the dimer (Figure 15b). Considering this, we believe that these NMR peaks originated from the inter-subunit interaction region of GAPDH. As GAPDH interacts with various biomolecules as cells age, it possibly interacts with them through interfaces exposed upon subunit dissociation, thereby acquiring moonlighting functions unrelated to glycolysis. To verify the hypothesis, we are examining O/P and O/R dimer variants, where the subunit labels O, P, Q, and R follow the nomenclature used in the PDB structure 1u8f (Figure 1a).

### Nitrosylation of the active site Cys residue alone does not directly cause major conformational changes, but oxidation of other residues likely leads to structural changes, such as denaturation

Gel filtration and NMR analyses of hGAPDH treated with CysNO showed that local nitrosylation of the active site residue Cys152 alone did not cause major structural changes (Figure 12). However, nitrosylation greatly reduced the thermal stability, suggesting that nitrosylation makes GAPDH more susceptible to oxidation. Furthermore, nitrosylation of Cys152 promoted the removal of NAD^+^ and brought GAPDH closer to its *apo* form. Oxidation involving nitrosylation of GAPDH with NOR3, which is more potent than CysNO, caused significant changes in the NMR spectra (Figure 14a,b). The broadening and disappearance of NMR peaks were possibly due to the denaturation of GAPDH or exchanges between multiple intra- and intermolecular conformations. Use of the C152S variant, in which local oxidation of the active site did not occur, resulted in decreased broadening of the NMR peaks (Figure 14c,d). Therefore, if oxidation and nitrosylation of Cys152 lead to significant conformational changes in GAPDH, this may be indirect, and there may be other mechanisms involved. This is consistent with a report by Samson *et al*. ^27^, who identified Met46 as the residue responsible for causing degeneration via oxidation. In our preliminary experiments, M46L variant was slightly more unstable than the wt, and M46I variant was too unstable to purify. These findings suggest the presence of other residues, in addition to Cys152, whose oxidation causes enzyme denaturation. Moreover, modification of Met46 significantly affects the overall structure of GAPDH as the residue is packed at the interface between subunits O and R ^50^.

### Currently, low-temperature conditions are optimal for detecting the subunit dissociated state

*apo* GAPDHs from pigs ^16,31^, rabbits, and yeast undergo subunit dissociation at low temperatures, possibly due to cold denaturation ^51^, in which hydrophobic interactions weaken at low temperatures. As high salt concentrations also promote GAPDH dissociation, both electrostatic and hydrophobic interactions are involved at subunit interfaces ^3^. However, at typical mammalian body temperatures, as shown by our NMR analyses at 40 °C, structural changes are limited to a small molar fraction of GAPDH. ESI-MS analysis of *apo* hGAPDH at room temperature revealed only its homotetrameric form. Nevertheless, as GAPDH is an abundant protein, accounting for approximately 10% of the total soluble proteins in cells, even a small molar ratio of transformed GAPDH is sufficient to evoke cellular moonlighting functions, such as binding to Siah1, migrating to the nucleus, and ultimately inducing cell apoptosis. Therefore, low temperatures can aid in detecting possible structural changes in GAPDH and in assessing the interactions involved in its moonlighting functions.

### Conclusion

In this study, we prepared pGAPDH and hGAPDH using an *E. coli* expression system and analyzed their conformations under various conditions. Unlike previous reports, subunit dissociation did not occur in the *apo* state, even at low temperatures and concentrations, as long as the solution was under reducing conditions. In contrast, when ATP or NOR3 was added, dimers or monomers were formed, and NMR signals suggesting flexible hydrophobic residues were observed. NAD^+^ binding significantly stabilized GAPDH in its homotetrameric state. Notably, thermostability of pGAPDH was always lower than that of hGAPDH under the tested conditions; however, various characteristics, such as subunit cleavage, were similar between pGAPDH and hGAPDH. Moreover, the local nitrosylation of Cys152 at the active site alone did not cause a significant conformational change. However, major conformational changes were observed when the unstable *apo* form, which is susceptible to oxidative attack, was oxidized at multiple sites. We are currently assigning NMR ^1^H/^13^C signals of methyl groups to identify the interaction sites with Siah1 ^7^, α-synuclein ^20^, amyloid-β ^39^, LDH ^41^, MDH1 ^43^, and cytokine mRNAs ^45^. The structures of these complexes are possibly dynamic and transient, warranting further NMR analyses.

## Materials and Methods

### Expression of isotopically labeled GAPDHs

DNAs encoding hGAPDH and pGAPDH were synthesized (GenScript) and introduced into pET47b plasmids (Merck) between the recognition sites of the restriction enzymes, *SmaI* and *SacI*, at a multicloning site. A (His)_6_-tag was fused to the N-terminus of the expressed protein. Removal of the tag by HRV3C PreScission protease left Gly-Pro-Gly at the N-terminus of GAPDHs.

Uniformly ^13^C/^15^N-labeled hGAPDH and pGAPDH were expressed in *E.coli* BL21(DE3) cells cultured in 1 L M9 minimal medium supplemented with 2 g/L [^13^C_6_]-glucose, 2 g/L ^15^NH_4_Cl, and 50 µg/mL kanamycin. *E.coli* cells for the expression of {*u*-[^2^H, ^15^N]; Ile^δ1^-[^13^CH_3_]; Met^ε^-[^13^CH_3_]; Leu^δ^, Val^γ^-[^13^CH_3_, ^12^CD_3_]}-hGAPDH were cultured in 100 mL D_2_O-based M9 minimal medium containing 2 g/L [^2^H]-glucose and 2 g/L ^15^NH_4_Cl. When the optical density at 600 nm (OD_600_) reached 0.5, the medium was supplemented with 200 mg/L α-ketobutyrate ([3,3-^2^H_2_, 4-^13^C_1_] 2-oxobutanoic acid), 200 mg/L α-ketoisovalerate (3-[^13^C_1_]methyl-[3,4,4,4-^2^H_4_] 2-oxobutanoic acid), and 40 mg/L [^13^C^ε^]-Met, and cooled to 15 °C ^19^. After 1 h, expression was induced with 50 µM isopropyl β-D-thiogalactopyranoside (IPTG), and after 14 h, *E. coli* cells were collected via centrifugation (6,000 rpm) at 4 °C for 30 min.

### Purification of GAPDHs

Cells were suspended in buffer A (20 mM Tris-HCl [pH 8.0], 400 mM NaCl, 40 mM imidazole, 1 mM DTT, 10 µM AEBSF, and 1% NP-40) and disrupted via sonication. The supernatant obtained after centrifugation (15,000 rpm) at 4 °C for 30 min was passed through the DEAE-Sepharose column to remove the nucleic acids and applied to the Ni-NTA Sepharose column (HisTrap FF crude, Cytiva). After washing the column with Tris-HCl buffer (pH 8.0) containing 40 mM imidazole and 10 or 400 mM NaCl, the proteins were eluted with buffer B (20 mM Tris-HCl [pH 8.0], 10 or 400 mM NaCl, 400 mM imidazole, and 1 mM DTT).

The solvent was exchanged with buffer C (20 mM Tris-HCl [pH 8.0] or 20 mM sodium phosphate [pH 7.4], 150 mM NaCl, 1 mM DTT, and 1 mM ethylenediaminetetraacetic acid (EDTA)) using an ultra-centrifugal filter (Amicon ultra-4 [30 kDa]; Merck). The (His)_6_-tag was cleaved by reacting with HRV3C overnight at 4 °C. The reaction solution was then passed through the Ni-NTA Sepharose column to remove the tags and uncleaved proteins. Proteins in the flow-through fraction were concentrated and fractionated on a size-exclusion column (HiLoad 16/60 Superdex 200pg) equilibrated with buffer C at a flow rate of 1 mL/min. Purified GAPDH was treated with activated charcoal (Extra Pure Reagent treated with HCl; Nacalai) (approximately 30 µg per 300 µL solution) on ice for 15 min, and the amount of bound NAD^+^ was determined by estimating the absorbance ratio (A_260_/A_280_) ^16^. When the value was < 0.6, activated charcoal was removed using a 0.22-µm filter. If the absorbance ratio did not reach the target value, the GAPDH solution was passed through the DEAE column with 150 mM NaCl, and activated charcoal treatment was repeated.

### Nitrosylation with CysNO

CysNO was prepared as follows: 5 mmol cysteine (0.61 g) was dissolved in 6 mL of 1 M HCl. Then, 5 mmol sodium nitrite (0.35 g of NaNO_2_) was dissolved in 1 mL water. The solutions were mixed and incubated in the dark for 30 min at room temperature. After centrifugation, pH of the supernatant was adjusted to 7.4 with 10 M NaOH. The amount of CysNO obtained was estimated by measuring the absorbance at 540 nm using S-nitrosoglutathione (Dojindo) as a reference (Saville–Griess assay) ^52^. Nitrosylated GAPDH was prepared by reacting purified GAPDH with 50-fold molar equivalents of CysNO for 60 min in the dark at room temperature and removing the excess CysNO via ultrafiltration. Modified and unmodified GAPDHs were denatured in aqueous solutions containing 0.1% formic acid and 50% acetonitrile and subjected to ESI-MS.

### Nitrosylation and oxidation with NOR3

GAPDHs were reacted with NOR3 dissolved in *d*-DMSO. A stock solution of NOR3 was added to GAPDH to create a mixed solution containing 4 µM deuterated GAPDH, 100 µM NOR3, and 1% *d*-DMSO.

### NMR measurements

NMR spectra were acquired at various temperatures (5–40 °C) using Bruker BioSpin Avance III HD spectrometers at ^1^H resonance frequencies of 500, 800, and 950 MHz with TCI triple-resonance cryogenic probes. For [^13^C, ^15^N]-GAPDHs, 2D ^1^H-^13^C HSQC echo-antiecho spectra were measured. For selectively and isotopically labeled GAPDHs with deuterium ([^2^H]), nitrogen-15 ([^15^N]), and specific methyl groups enriched with carbon-13 ([^13^C^1^H_3_]) and deuterium ([^12^CD_3_]), 2D ^1^H-^13^C methyl-TROSY HMQC spectra were measured ^21^. The typical spectral widths were 24 ppm (^1^H) and 50 ppm (^13^C), with 2,048 and 200 data points for HSQC, respectively. For TROSY-HMQC, the widths were 17 ppm (^1^H) and 30 ppm (^13^C), with 2,048 and 300 data points, respectively. The other measurement conditions, including protein concentration, were slightly different depending on the specific experiments performed and are described in each figure legend.

### ESI-MS measurements

Solvents containing 5–10 µM GAPDH were replaced with 10 mM ammonium acetate buffer using an ultra-centrifugal filter (30 kDa cut-off), and the sample was subjected to ESI-MS under native-like conditions. For mass determination of nitrosylated GAPDH, the sample was denatured with 0.1% formic acid and 50% acetonitrile, and subjected to ESI-MS (Synapt G2 HDMS, Waters). Subsequently, 4 µL of the sample was injected using the Pt-coated glass capillary (HUMANIX).

### Thermal shift assay

Protein solution was prepared at a concentration of 0.1 mg/mL. The fluorescent dye (SYPRO Orange Protein Gel Stain) was diluted 50-fold with a buffer and added to the protein solution at 1/20 dilution. Subsequently, 20 µL of the prepared sample was dispensed into a 96-well plate.

The fluorescence emitted was measured using the C1000 Thermal Cycler (Bio-Rad). After equilibration at 25 °C for 1 min, the temperature was increased to 90 °C in 0.5 °C/10 s increments to plot the melting curve.

### Analytical gel filtration chromatography

Samples were applied to a column (Superdex 200 10/300GL) using a pump (AKTA prime) at a flow rate of 0.3–0.6 mL/min.

### Measurement of specific activity of GAPDH

Activity was estimated by measuring the absorbance of NADH in real-time at 340 nm (extinction coefficient of 6, 220/M/cm). To 325 µL of enzyme solution (40 mM Tris-HCl [pH 8.0], 25 mM Na_2_HAsO_4_, 1 mM NAD^+^, 1 mM EDTA, 10% CHAPS, and 0.01 µM GAPDH subunit), 125 µL of 4 mM glyceraldehyde-3-phosphate was added to make the final concentration of the substrate 1 mM. The mixture was immediately vortexed and transferred to a quartz cuvette for measurements. Measurements were taken every 5 s for 20 min. The conversion from glyceraldehyde-3-phosphate to 1-arseno-3-phosphoglycerate occurred in the enzyme solution. The reverse enzymatic reaction was suppressed as 1-arseno-3-phosphoglycerate spontaneously degraded to 3-phosphoglyceric acid via hydrolysis. The specific activity of the enzyme (U µmol/mg/min) was calculated from the slope at 0 min using a graph showing the absorbance at 340 nm vs. reaction time.

### Preparation of *α*-synuclein

*E. coli* BL21(DE3) cells were transformed with plasmids containing the α-synuclein gene and cultured in 1 L of M9 medium containing ^15^NH_4_Cl and [^13^C]-glucose at 37 °C. When the OD_600_ reached 0.6, IPTG was added to a final concentration of 0.1 mM, and the cells were further cultured at 15 °C for 5 h. The cells, collected by centrifugation, were disrupted by sonication, and the supernatant was applied to a DEAE column (50 mM Tris-HCl, pH 7.2, 400 mM NaCl) to remove nucleic acids. The solution was heat-treated at 80–85 °C for 15 min. The supernatant was then applied to a Q column (50 mM phosphate buffer, pH 6.5), and α-synuclein was eluted using a NaCl concentration gradient of up to 1 M. Finally, α-synuclein was purified using gel filtration chromatography (HiLoad 16/60 Superdex 200pg).

### Preparation of Siah1

The gene encoding full-length Siah1 (282 residues) was subcloned into the pCold vector. The expressed protein contained an N-terminal (His)_6_-tag, trigger-factor, and the TEV protease recognition sequence. The purification method was essentially the same as that used for other (His)_6_-tagged proteins mentioned earlier. After elution from the Ni-NTA column, Siah1 was separated from the N-terminal portion through cleavage by the TEV protease. Since SDS-PAGE analysis suggested a nonspecific interaction between trigger-factor and Siah1, the two were separated by washing the Ni-NTA column with a buffer containing 1 M urea.

### Preparation of mRNA-containing samples

Single-stranded RNAs consisting of 9, 13, and 38 nucleotides were synthesized by Fasmac. Each RNA was mixed with [^15^N, ^13^C]-hGAPDH. To prevent RNA degradation, NMR sample tubes were soaked in 0.1 M NaOH for 1 h, and 400 units of Recombinant RNase Inhibitor (Takara Bio Inc.) were added to the samples. The RNA sequences were as follows.

38-nt RNA: 5’-GUG AUU AUU UAU UAU UUA UUU AUU AUU UAU UUA UUU AG-3’ 13-nt RNA: 5’-AUU UAU UUA UUU A-3’ 9-nt RNA: 5’-AUU UAU UUA-3’

## Author Contributions

T.K., S.A., H.A., and T.I. designed the experiments; H.S., Y.M., Y.W., M.T., K.A., and Y.M. prepared the samples and performed the experiments; T.I. wrote the paper.

## Funding

This study was funded by JSPS KAKENHI (Grant-in-Aid for Scientific Research (C); grant number 23K05667), Grant-in-Aid for Transformative Research Areas (A) (grant number 2323H04958), Basis for Supporting Innovative Drug Discovery and Life Science Research (BINDS; grant number JP23ama121001), and a grant for academic research from Yokohama City University.

## Acknowledgements

T.I. is grateful to the following students for their contributions to this research: Satomi Komiya, Kaede Hamada, Mizuho Komatsu, Mahoko Harada, Yumi Suzuki, Ayaka Kondo, Tomoya Takaira, Kento Murase, Kouhei Arai, Shuhei Yamaura, and Shuri Sakuma. Additionally, T.I. would like to thank Dr. Aritaka Nagadoi for his guidance and insightful discussions throughout this project. We would like to thank Editage (www.editage.jp) for English language editing. This study used the NMRbox cloud system at the National Center for Biomolecular NMR Data Processing and Analysis (BTRR).

## Conflicts of Interest

The authors declare no conflicts of interest.

## Notes

### Competing Interest Statement

The authors have declared no competing interest.

